# Surface-based tracking for short association fibre tractography

**DOI:** 10.1101/2021.05.07.443084

**Authors:** Dmitri Shastin, Sila Genc, Greg D. Parker, Kristin Koller, Chantal M.W. Tax, John Evans, Khalid Hamandi, William P. Gray, Derek K. Jones, Maxime Chamberland

**Author notes:** Corresponding author. Address: CUBRIC, Maindy Rd, Cardiff, United Kingdom, CF24 4HQ *Email address* (Dmitri Shastin).

## Abstract

It is estimated that in the human brain, short association fibres (SAF) represent more than half of the total white matter volume and their involvement has been implicated in a range of neurological and psychiatric conditions. This population of fibres, however, remains relatively understudied in the neuroimaging literature. Some of the challenges pertinent to the mapping of SAF include their variable anatomical course and proximity to the cortical mantle, leading to partial volume effects and potentially affecting streamline trajectory estimation. This work considers the impact of seeding and filtering strategies and choice of scanner, acquisition, data resampling to propose a whole-brain, surface-based short (≤30-40 mm) SAF tractography approach. The framework is shown to produce longer streamlines with a predilection for connecting gyri as well as high cortical coverage. We further demonstrate that certain areas of subcortical white matter become disproportionally underrepresented in diffusion-weighted MRI data with lower angular and spatial resolution and weaker diffusion weighting; however, collecting data with stronger gradients than are usually available clinically has minimal impact, making our framework translatable to data collected on commonly available hardware. Finally, the tractograms are examined using voxel- and surface-based measures of consistency, demonstrating moderate reliability, low repeatability and high between-subject variability, urging caution when streamline count-based analyses of SAF are performed.

## 1. Introduction

Functional integration of the brain subunits is mediated in part by the white matter (Neubert et al. (2010)) which comprises a vast network of connections between neuronal populations and has been shown to exhibit change in response to physiological processes (Scholz et al. (2009); Hihara et al. (2006); Dubois et al. (2014); de Groot et al. (2015); Slater et al. (2019)) and disease (Mito et al. (2018); Datta et al. (2017); de Schipper et al. (2019)). The white matter is typically divided into projection, commissural and association fibres. It is estimated that the association fibres, connecting the cortical areas within hemispheres, dominate the white matter (Schüz and Braitenberg (2002)). They are in turn subdivided into long and short range (local) fibres, sometimes also distinguishing neighbourhood association fibres (Schmahmann and Pandya (2006)). The long-range fibres course in the depth of the white matter, connecting distant areas of the hemisphere and forming distinct bundles that have largely consistent anatomy across individuals. Conversely, the short association fibres (SAF) connect adjacent cortical areas. Their most superficial component is often referred to as ‘U-fibres’ and described as a thin band that runs immediately beneath the sixth layer of the cortex (Schmahmann and Pandya (2006)) encompassing a single gyrus or sulcus (Schüz and Braitenberg (2002)). It is established that neighbouring cortical areas exhibit the strongest structural connectivity (Markov et al. (2014)). Further, it is estimated that only □10% of the cortico-cortical connections belong to the long fascicles, with the volume of the U-fibres possibly as much as □60% of the total white matter volume (Schüz and Braitenberg (2002)). It is remarkable therefore that in the neuroimaging literature SAF have only started to gain attention recently (Ouyang et al. (2017)).

Diffusion MRI (dMRI) is the preferred method for studying structural properties and connectivity of white matter pathways *in vivo*. Its sensitivity to the random microscopic motion of water molecules (Stejskal and Tanner (1965)) enables judgements to be made regarding the local orientational architecture and microstructural properties of the fibres (Pierpaoli et al. (1996)). In the past few years, a number of dMRI-based studies have shown that SAF are affected by age and sex (Phillips et al. (2013)) as well as pathology including autism (d’Albis et al. (2018)), schizophrenia (Phillips et al. (2011), encephalitis (Phillips et al. (2018)) and epilepsy (O’Halloran et al. (2017); Liu et al. (2016); Govindan et al. (2013)), among others. dMRI methods used to study SAF can be broadly divided into those that do not use tractography and those that do. The former typically sample measures of microstructure in the superficial white matter as defined by regions of interest (Nazeri et al. (2013)) or uniformly along the cortical surface (Phillips et al. (2013, 2018, 2011); Liu et al. (2016)). This approach avoids any potential biases of tractography and can be less affected by the differences in cortical folding by using surface registration (Fischl et al. (1999)) but it is insensitive to the thickness and shape of the cortico-cortical bundles. On the other hand, tractography-based methods capitalise on fibre orientation modelling which allows reconstruction of streamlines providing information about white matter morphology (Mori and Van Zijl (2002)). Recent advances in image acquisition (Jones et al. (2018)) and processing as well as development of advanced fibre orientation estimation methods (Tournier et al. (2007); Dhollander et al. (2016); Jeurissen et al. (2014)) and streamline integration and filtering algorithms (Smith et al. (2012); Daducci et al. (2015); Smith et al. (2015)) have improved the quality of tractography. Despite this, available tools are typically used to study whole-brain tractograms or focus on the deep white matter bundles that show consistent organisation across individuals and thus the performance of these tools for investigating SAF remains uncertain.

## 2. Challenges in SAF reconstruction

The study of SAF is confounded by a number of anatomical considerations and methodological limitations (for an overview, see Guevara et al. (2020); Jeurissen et al. (2017); Rheault et al. (2020); Reveley et al. (2015)) which span initial tractogram generation, SAF-specific filtering and analysis.

### 2.1. Tractogram generation

Tractogram generation faces the challenges of partial volume effects (due to the proximity of SAF to the cortex and CSF spaces) and complex local anatomy with crossing, bending, kissing, and fanning fibres. Their subcortical course makes SAF potentially more sensitive to the so-called “gyral bias” – the phenomenon in tractography where many more streamlines terminate in the gyral crowns as opposed to the sulcal fundi (Li et al. (2010); Nie et al. (2011); Chen et al. (2012); Van Essen et al. (2014); Schilling et al. (2018), Cottaar et al. (2021)). This is disproportionate in relation to the respective axonal connections (which also favour the gyri) as reconstruction algorithms fail to produce the sharp turn towards the gyral wall seen on histology (Schilling et al. (2018)). Specifically in the context of U-fibres, the gyral bias has been demonstrated even when state-of-the-art acquisition and tractography were used (Movahedian Attar et al. (2020)). Additionally, streamline reconstruction can experience difficulty traversing the subcortical white matter preferring a tangential trajectory instead (Reveley et al. (2015)). This tendency is highly undesirable for the long-range connections but may prove beneficial for SAF as tracking is encouraged along the natural SAF course. It is therefore important to investigate the influence of these effects on SAF tractography.

### 2.2. Tractogram filtering

From the spatial filtering perspective, SAF may be defined locally based on manual dissections (Catani et al. (2012), Catani et al. (2017)) or functional MRI signal-derived cortical regions of interest (Movahedian Attar et al. (2020)), whilst for globally (brain-wise) defined SAF, the filtering criteria typically involve size, shape/location and/or cortical parcellation. Despite the existence of studies examining the anatomical course of SAF in isolated brain regions (Catani et al. (2012), Catani et al. (2017)), the absence of a detailed anatomical knowledge regarding the distribution and consistency of SAF on a whole-brain level or even a universally accepted definition (Ouyang et al. (2017)) complicates development and validation of non-invasive methods dedicated to the study of this subset of the white matter. For instance, the length definition of SAF (or U-fibres) varies across sources. Some authors have focused on the relatively long streamlines of 20-80 mm (Guevara et al. (2017); Kai and Khan (2019)) or more (Román et al. (2017)), mainly concerning the bundles connecting neighbouring gyri; while others (Song et al. (2014); Movahedian Attar et al. (2020)) included the smaller range of 3-30 mm based on the classification by Schüz and Braitenberg (2002). Next, although using streamline similarity measures (typically shape and distance metrics) as filtering criteria (Román et al. (2017); O’Halloran et al. (2017); Kai and Khan (2019)) may appear appealing, this may lead to exclusion of otherwise valid streamlines as SAF have been demonstrated to exhibit complex, diverse morphology (Movahedian Attar et al. (2020)) and varying spatial overlap (Zhang et al. (2010)); this is particularly true for shorter (<35 mm) streamlines (Román et al. (2017)). The use of cortical parcellations (division of the cortical mantle into discrete areas) can carry uncertainties of its own. The choice of parcellation scheme, termination criteria during tracking, and the way streamlines are associated with individual parcels all influence the result (Yeh et al. (2019)).

### 2.3. Tractogram comparison

Group-wise analysis of SAF is challenged by inter-subject variation in cortical folding (Rademacher (2002)). Even the sulci known to exhibit more anatomical consistency across individuals (such as those corresponding to the primary somatosensory areas (Rademacher (2002)) demonstrate individual morphological differences up to 1-2 cm in a common reference frame (Steinmetz et al. (1989)). The trajectories of short (up to 40 mm) superficial streamlines appear to be strongly influenced by the gyral pattern (Bajada et al. (2019)). Taken together, one should expect low consistency when comparing SAF tractograms composed of shorter streamlines between individuals based on their shape or spatial distribution alone. Connectome-based comparisons using cortical parcellations are possible yet again they face the same challenges as described above.

The aim of the current study was to develop a methodological framework suitable for whole-brain tractography of SAF ≤30-40 mm. To produce SAF-specific tractograms, we introduced simple, anatomy-driven filtering criteria that did not require manual dissection/pruning or the use of additional shape/parcellation-based priors. We hypothesised that distributing streamline seeds directly on the white matter surface as well as employing surface-based filtering techniques will be less prone to discretisation errors and promote more desirable streamline features in regions with fine anatomical detail. To this end, we defined a set of descriptive features and used them to compare surface-based tractograms against tractograms generated with a voxel-based method. We followed this with an investigation into how these features changed depending on angular and spatial resolution, number of shells and b-values by comparing different diffusion acquisitions, as well as performing dMRI data interpolation to smaller or larger voxels. The tractograms were then assessed for within-subject consistency to evaluate the framework, coupled with between-subject analyses to inform the reader of what could be expected during the use of the method for population studies.

## 3. Material and methods

### 3.1. Proposed surface-based SAF tractography framework

The overall workflow is summarised in Figure 1. A FreeSurfer-generated fine cortical mesh (Fischl (2012)) is used to place streamline seeds in dMRI space. Tracking is performed using a probabilistic tracking algorithm (described below) with fibre orientation distribution functions (fODFs) generated from multishell dMRI data (Table 2).

**Figure 1:**
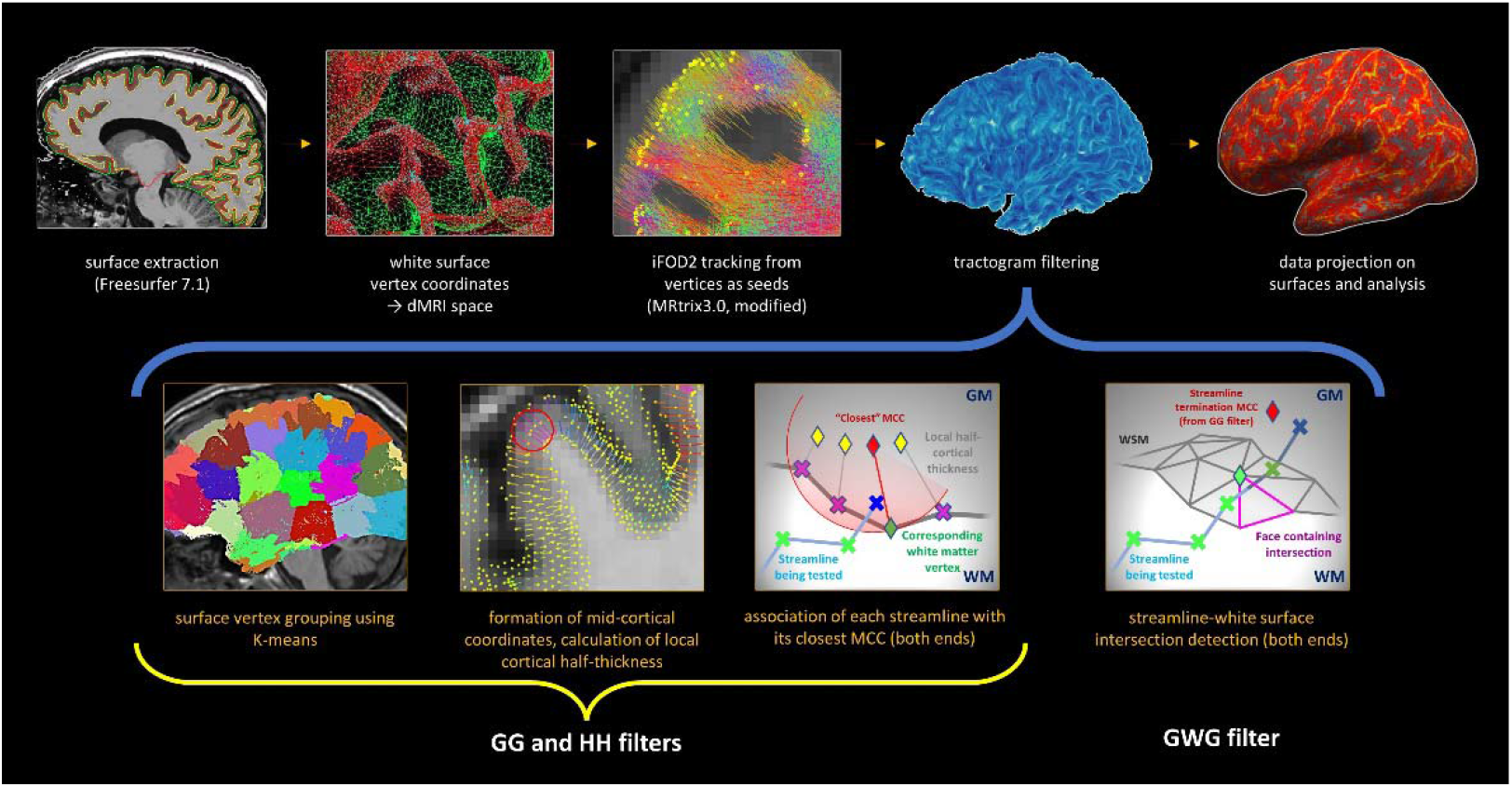
Pipeline summary. After seeding from white surface mesh (WSM) coordinates (top row, 1-3), tractograms are filtered (top row, 4) to ensure each streamline starts and ends in the neocortex (grey-grey filter) of the same hemisphere (hemisphere-hemisphere filter) and escapes into white matter along the way (grey-white-grey filter). The grey-grey filter (bottom row, 1-3) functions by finding the closest midcortical coordinate (MCC, average of the matching WSM and pial coordinates) for each streamline end (with K-means clustering of MCCs for speed - bottom row, 1). A streamline end is considered in the grey matter if it lays within the local cortical half-thickness of its MCC (bottom row, 3). Then, two intersections with WSM (one either end) are sought (bottom row, 4) at which point the intracortical portion is truncated. The optional surface-based analysis is conducted after the filtering (top row, 5).

**Table 1:** List of abbreviations

| Abbreviation | Meaning/Interpretation |
| --- | --- |
| ACT | Anatomically-constrained tractography |
| CVB | Between-subject coefficient of variation |
| CVW | Within-subject coefficient of variation |
| dMRI | Diffusion magnetic resonance imaging |
| DSI | Diffusion spectrum imaging |
| DTI | Diffusion tensor imaging |
| FA | Fractional anisotropy |
| fODF | Fibre orientation distribution function |
| FWHM | Full-width half-maximum |
| GG | Grey-grey filter |
| GMWMI | Grey matter – white matter interface |
| GWG | Grey-white-grey filter |
| HARDI | High angular resolution diffusion imaging |
| HH | Hemisphere-hemisphere filter |
| ICC | Intraclass correlation coefficient |
| LCHT | Local cortical half-thickness |
| MGA | Maximum gradient amplitude |
| MCC | Mid-cortical coordinate |
| SA | “State-of-the-art” acquisition |
| SAF | Short association fibres |
| ST | “Standard” acquisition |
| TDI | Track density imaging |
| WSM | White surface mesh |

**Table 2:**
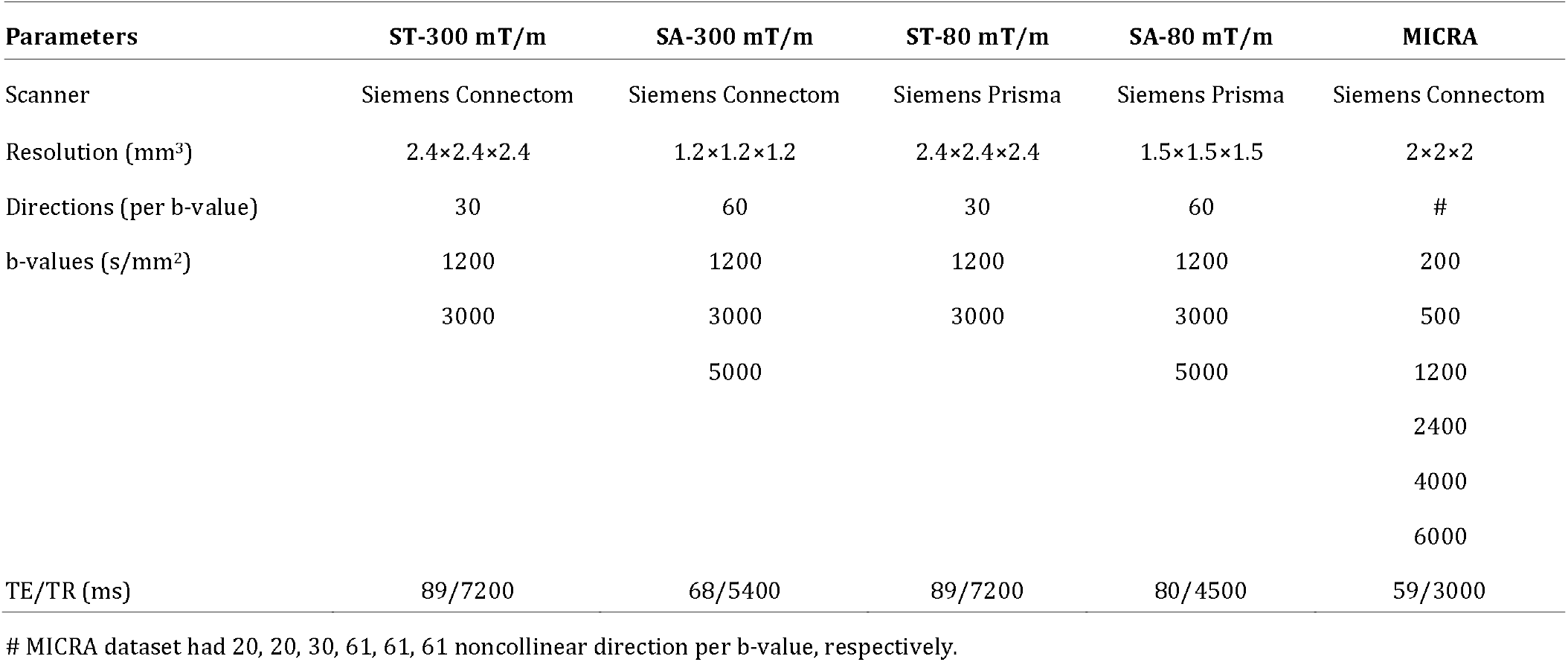
Acquisition parameters utilised for dMRI sequences in the two cohorts studied. TE, echo time. TR, repetition time.

| Parameters | ST-300 mT/m | SA-300 mT/m | ST-80 mT/m | SA-80 mT/m | MICRA |
| --- | --- | --- | --- | --- | --- |
| Scanner | Siemens Connectom | Siemens Connectom | Siemens Prisma | Siemens Prisma | Siemens Connectom |
| Resolution (mm<sup>3</sup>) | 2.4×2.4×2.4 | 1.2×1.2×1.2 | 2.4×2.4×2.4 | 1.5×1.5×1.5 | 2×2×2 |
| Directions (per b-value) | 30 | 60 | 30 | 60 | # |
| b-values (s/mm<sup>2</sup>) | 1200 | 1200 | 1200 | 1200 | 200 |
|  | 3000 | 3000 | 3000 | 3000 | 500 |
|  |  | 5000 |  | 5000 | 1200 |
|  |  |  |  |  | 2400 |
|  |  |  |  |  | 4000 |
|  |  |  |  |  | 6000 |
| TE/TR (ms) | 89/7200 | 68/5400 | 89/7200 | 80/4500 | 59/3000 |

By definition, a streamline representing a population of association fibres must satisfy the following criteria: (1) both ends terminate in the neocortex; (2) both ends terminate in the same hemisphere; (3) the streamline courses through the white matter. Three respective filters ensuring these three criteria are met are applied to the initial tractogram. First, for each streamline end the closest point on the mid-cortical mesh is identified. The streamline terminates within the cortex if the distance to this closest point is no greater than half of the cortical thickness at that point. Second, a prior knowledge of which hemisphere each surface point belongs to ensures that the streamline starts and ends in the same hemisphere. Finally, the course within the white matter is confirmed by identifying two intersections (one either end of the streamline) with the white matter surface. The following paragraphs detail each of the steps described.

#### 3.1.1. Initial streamline generation

The FreeSurfer white matter surface mesh (WSM) typically contains □1.5 vertices/mm^2^ for a total of □250K vertices (points) for both hemispheres (excluding the medial wall) and a face (triangle) area of ~ 0.3 mm^2^ (range: 0.07-0.7 mm^2^, top and bottom 2% excluded) representing a reasonably dense and even spread. This can be further remeshed if needed: in this work, triangle edges whose length exceeded the mean by two standard deviations are bisected (resulting in more normally distributed edge length and triangle area histograms) to generate additional seeding points. Point coordinates are then transformed to dMRI space with ANTs (Avants et al. (2009)) using the inverse warp (see subsection 3.2.1 for registration details) and concatenated into a single array used to initiate seeding with MRtrix 3.0 (Tournier et al. (2019)). To this end, MRtrix was modified such that it could read coordinates from the array and use them as seeds with equal weights during tractogram generation. This seeding mechanism was verified by visual inspection of seed distribution on T1-weighted volumes in dMRI space and by comparing input and output seed coordinates (data not shown). Next, tracking is performed using the second-order integration probabilistic algorithm iFOD2 (Tournier et al. (2010)) which can handle the expected large number of fibre crossings and challenging morphology; probabilistic tracking has previously been demonstrated to result in less gyral bias (Nie et al. (2011)), potential for better performance on the receiver-operator characteristic curve (Grisot et al. (2021)) and greater spatial overlap of SAF (Guevara et al. (2020)). The “select” parameter (number of streamlines in the unfiltered tractogram) is set to 5 million to ensure an adequate number of streamlines per vertex. An additional restriction on the maximum streamline length of 40 mm is used to be consistent with the SAF definition of Schüz and Braitenberg (2002) also accounting for the fact that intracortical portions are later truncated (see section 3.1.2). Importantly, the fODF amplitude threshold is kept sufficiently low at 0.05 to facilitate tracking at the grey-white interface. Other parameters are left at default settings in the MRtrix implementation of iFOD2 (max angle: 45°, step size: 0.5 mm, fODF power: 0.25).

#### 3.1.2. Streamline filtering

##### Grey-grey (GG) filter

To identify streamlines starting and ending in the neocortex, midcortical coordinates (MCC) are defined by averaging coordinates of the matching WSM and pial surface mesh vertices. Next, local cortical half-thickness (LCHT) is defined as the Euclidean distance between the MCC and the corresponding WSM vertex to account for local variation in cortical thickness. Both ends of each streamline in the initial tractogram are then evaluated for “intracortical position” by (1) identifying the closest MCC, (2) measuring the Euclidean distance to it, (3) comparing this distance to the LCHT measure of said MCC. The “intracortical position” is confirmed if the streamline terminates in a sphere centred on the MCC and with the radius LCHT (see Appendix A).

To improve computational efficiency, all MCCs are clustered with K-means using squared Euclidean distances (Arthur and Vassilvitskii (2006)); finding the centroid closest to a streamline end means the closest MCC needs only be identified within the cluster of that centroid (see Appendix B).

##### Hemisphere-hemisphere (HH) filter

The original hemispheric membership (left or right) of all cortical vertices and thus MCCs is known; the hemispheric allocation of each streamline end is then straightforward once its closest MCC is identified during GG filtering. This filter acts by only preserving streamlines whose both ends reside in the same hemisphere.

##### Grey-white-grey (GWG) filter

After ensuring all streamlines terminate

*for each hemisphere:*
*for each streamline:*
*define the bounding box*
*find all surface faces within the bounding box*
*define a bounding box for each identified face*
*define a bounding box for each individual streamline segment*
*register intersection between face and segment bounding boxes*
*for each bounding box intersection:*
*check for segment-triangle intersection*

intracortically, the last filter needs only to detect escape into white matter. Streamlines travel some distance within the cortex before this happens (mean: 3-4 mm, median: 2-3 mm), typically resulting in a non-correspondence between the MCC associated with a streamline’s end and the streamline’s intersection with WSM. Due to the possibility of this occurring on a subvoxel scale, and because the exact point of intersection with WSM is of interest, filtering is performed by detecting intersections between streamline segments and WSM faces instead of applying a white matter mask. As considering all segments of all streamlines with all WSM faces would be extremely inefficient, intersections between bounding boxes are used in the initial step to significantly restrict the search space. The pseudocode is provided below:

Segment-triangle intersection detection is performed using signed volumes but as this is approximately three times slower compared to matching segment and face bounding boxes, the latter is done first. In addition to detecting escape into white matter, the filter allows to associate each streamline end with its nearest WSM vertex and truncate streamlines at that point if desired (enabled in this work). Importantly, in the proposed framework we assume that streamlines generally follow the grey-white interface and ignore any WSM intersections that occur between the two intracortical ends (the consequences of that are examined in Results).

### 3.2. Framework evaluation

Evaluation took a multi-phase approach. First, the results from the proposed surface-based framework were compared against a voxel-based method and the differences explored. Next, the roles of scanner, acquisition and dMRI data resampling on framework performance were evaluated for parameter optimisation. Finally, experiments were carried out to evaluate within- and between-subject consistency of SAF tractograms obtained using the proposed framework.

#### 3.2.1. Data acquisition and pre-processing

Acquisition parameters for all protocols are summarised in Table 2.

##### Cohort A

Repeatability data from the MICRA study (Koller et al. (2020)) were used to assess the different methods (subsection 3.2.2) and the consistency of results (subsection 3.2.4). In short, after a written informed consent, brain MR data of six healthy adults (3 males and 3 females, age range 24-30) were obtained using the Siemens Connectom (MGA=300 mT/m) scanner. Each participant was imaged five times using the same protocol within a two-week period at approximately the same time of day.

##### Cohort B

Tractograms were evaluated using a data set of 14 subjects (4 males and 10 females, age range 16-30) acquired on two different 3T scanners with different maximum gradient amplitudes (MGA): Siemens Connectom (MGA=300 mT/m) and Siemens Prisma (MGA=80 mT/m) (Tax et al. (2019)). Two acquisition protocols were used: “standard” (ST) and “state-of-the-art” (SA), with the latter having higher spatial and angular resolution achieved with multiband acquisition and stronger gradients to shorten echo time (Table 2). This, in turn, enabled a higher signal-to-noise ratio per unit time for a given b-value, allowing for utilisation of higher b-values which are more sensitive to intra-axonal water displacement (Jones et al. (2018); Setsompop et al. (2013); Genc et al. (2020)). Written informed consent was given by all subjects.

##### Pre-processing

Spin-echo echo-planar dMRI images were corrected for slice-wise intensity outliers (Sairanen et al. (2018)), signal drift (Vos et al. (2017)), Gibbs artifact (Kellner et al. (2016)), eddy current distortion and motion artifact (Andersson and Sotiropoulos (2016)), echo-planar image distortion (Andersson et al. (2003)), and gradient non-linearities (Glasser et al. (2013); Rudrapatna et al. (2018)). In all experiments, dMRI data resampling took place at the end of pre-processing and before tractogram generation. In all cases, data were upsampled to 1×1×1 mm^3^ (Dyrby et al. (2014)) and, for the evaluation of voxel size effects (Cohort B), resampled to 2×2×2 mm^3^. Diffusion tensor estimation in each voxel was performed with nonlinear least squares (Jones and Basser (2004)). The fODF (Tournier et al. (2007)) was derived using 3-tissue response function estimation (Dhollander et al. (2016)) and subsequent multi-shell multi-tissue constrained spherical decomposition (Jeurissen et al. (2014)) with harmonic fits up to the eighth order. The quality of the pre-processing steps as well as fODFs were visually confirmed for all subjects.

Anatomical data (Siemens MPRAGE1 sequence, voxel size: 1×1×1 mm^3^, TR/TE 2300/2.81 ms) were run through the FreeSurfer 7.1 package (Fischl (2012)) which includes standard T1-weighted volume pre-processing steps. For Cohort A, the longitudinal stream designed for repeated acquisitions was used (Reuter et al. (2012)). One subject lacked a T1-weighted volume for one of the sessions; instead, the within subject template (referred to as “base” in the longitudinal stream) was used, resulting in a total of six “base” and twenty-nine final (referred to as “long”) sets. The quality of produced surface meshes was visually inspected at every step and corrected where necessary as per the standard FreeSurfer protocol. dMRI-derived fractional anisotropy (FA) volumes were non-linearly registered to FreeSurfer T1-derived “brain” volumes using ANTs; coordinates of surface vertices were then brought into dMRI space using the inverse transform and registration quality was visually confirmed in each case. An average subject template was created for group analyses from the six “base” sets with FreeSurfer’s *make_average_subject* command and used it as a common space template for all “long” sets (surface co-registration done with *surfreg*).

#### 3.2.2. Tractogram evaluation

The following section describes assessments used in the first two experiments (comparison with voxel-based method and effects of scanner, acquisition, resampling). Depending on the experiment, some assessments were avoided if not relevant.

##### General characteristics

SAF tractograms were characterised with the following features:

For the purposes of “coverage”, “coverage bias”, “streamline ends in gyri”, mesh vertices with negative mean curvature were considered to lie in gyri and the rest in sulci. The difference between “streamline ends in gyri” and “coverage bias” was that for the latter the streamline numbers were not taken into account, and only parts of the cortical surface covered were considered.

##### Streamline angle at the cortex

Axonal projections follow a near-radial course in gyral crowns and a more tangential course along sulcal banks and fundi (Van Essen et al. (2014); Schilling et al. (2018)). To assess whether SAF tractograms were consistent with this pattern, position along the gyral blade was discretised into equally spaced bins and the average angle between the first streamline segment and the cortex (calculated as 90° □ angle to normal) per bin was plotted against the position on the gyral blade. The cortical surface was then subdivided according to position along the gyral blade (mean curvature values) into five zones of equal area (from gyral crowns to sulcal fundi) and the mean angle for each zone, as well as the overall mean angle, were calculated. Only the bins in the 5-95% range were used for this calculation as the bins at extreme ends had multiple empty values. Empty bins within the 5-95% range (few points in the gyral extremes) had values imputed using inverse distance weighted interpolation.

##### Distribution of connections along the gyral blade

Distribution of SAF streamlines in relation to gyri and sulci was further investigated by splitting position along the gyral blade into five regions of equal area (as in the previous paragraph) that were used as nodes. Undirected connectivity matrices were then constructed with edge weights determined by streamline counts similar to the recent work by Cottaar et al. (2021).

#### 3.2.3. Comparison with voxel-based method

##### Choice of benchmark and tractogram generation

It was hypothesised that voxel-based seeding and filtering are subject to discretisation errors resulting in uneven streamline end distribution, with smaller regions being misrepresented or excluded (Appendix C). To investigate how the proposed framework fared against voxel-based methods, the established MRtrix ACT/GMWMI was used for benchmarking.

To produce SAF tractograms, MRtrix ACT/GMWMI pipeline was modified in the following way. First, FreeSurfer’s “aseg” volume was transformed to dMRI space using the previously computed registration and nearest neighbour interpolation (preserving segmentation labels). This acted as the input into MRtrix’s FreeSurfer-based five-tissue-type (5TT) segmented tissue image generation algorithm (Smith et al. (2012)). The 5TT image was then manipulated such that the cerebellar cortex and the amygdala/hippocampus were excluded from the grey matter volume (matching the cortical areas used in surface seeding), while the deep nuclei as well as the ventricles were added to the white matter volume with their original volumes set to null. The manipulation effectively forced all streamlines to start and end at the neocortex. Following this, a grey matterwhite matter interface (GMWMI) volume was produced and tractogram generation performed with matching parameters (including the fODF threshold of 0.05 and the ≤40 mm length limit) until the total number of streamlines for ACT/GMWMI exceeded that of the surface method by 20%. Precise cropping at the grey-white matter interface was enabled. Then, for each streamline end the closest WSM vertex was found, and streamlines connecting the two hemispheres were discarded. Finally, streamline removal continued at random until the total number of streamlines equalled that of the surface method for each data set. Closest WSM vertices identified in the previous step allowed surface-based assessments of the ACT/GMWMI-produced streamlines analogously to those of the proposed framework.

#### 3.2.4. Effects of scanner, acquisition, resampling

##### Fibre orientation distribution functions

Scanner and sequence effects on fODF configuration were first evaluated visually. For this comparison only, all four data sets of a subject were registered to the same (subject-specific) T1-weighted volume, with fODFs computed prior to registration. Three separate depths were plotted for ease of comparison: grey-white interface, deep white matter, and cortex. The former was represented by WSM in the same space, while the latter two were formed by projecting the WSM coordinates by 1 mm deep or superficial, respectively, along the vectors of the matching WSM and pial surface vertices.

Next, in order to evaluate fODF orientation systematically, the first (largest) peaks of all fODFs in original subject dMRI space were extracted to represent the greatest probability of local streamline propagation. Peak orientation at the three depths described above was sampled for each vertex of the WSM using trilinear interpolation (mirroring the iFOD2 algorithm used for tractography) and the angle with surface normal was calculated using dot product. Angles at the three depths were then plotted against position on the gyral blade similar to the streamline-angle cortex plots described in “3.2.2. Tractogram evaluation, Streamline angle at the cortex”.

##### Tractogram comparison

Tractograms were brought into Prisma/ST/2 mm^3^ space with affine co-registration (12 degrees of freedom) of the corresponding FA volumes and subsequent transformation of streamline point coordinates using ANTs. After the tractograms were inspected visually, the effects of scanner, acquisition and resampling were additionally investigated using linear modelling with response variables as defined in “3.2.2. Tractogram evaluation, General characteristics”.

#### 3.2.5. Assessment of consistency

##### TDI maps

Volume-based track density imaging (TDI) maps for each session were generated with MRtrix’s *tckmap* command. The maps were compared in average subject space by applying the previously obtained dMRI-to-T1 transform followed by a “base-to-average” transform (concatenated and performed in a single step using ANTs).

##### Surface-based analysis

Each streamline end was associated with its closest WSM vertex allowing streamline-related metrics to be recorded at these vertices in subject anatomical space. This enabled the use of surface-based registration for comparisons, which due to the complexities of cortical folding, can be superior to volume-based registration for these superficial structures. The following measures on the surface were recorded: (1) number of streamlines per vertex; (2) mean streamline length per vertex; (3) mean FA per streamline per vertex. The latter can be thought of as sampling FA along the surface with a kernel whose size and shape changed informed by the streamlines at the vertex being sampled. For all surface-based analyses, only the cortical surface (excluding the “medial wall” label) was studied. Individual subjects’ results (Appendix D) were co-registered, stacked(*mris_preproc*) and smoothed (*mri_surf2surf*) at 5 mm full-width at half-maximum (FWHM). Smoothing is commonly used in neuroimaging to boost signal-to-noise ratio, alleviate registration misalignment, and improve normality of residuals (Jones et al. (2005)). As per-vertex testing was not sensitive to between-vertex interactions, smoothing provided an alternate means to account for these interactions.

#### 3.2.6. Statistical analysis

All statistical analyses were performed in MATLAB 2015a. Hypothesis testing was performed with a significance level of 0.05 in all cases.

##### Effects of surface seeding

Data were inspected and concluded to be approximately normally distributed (allowing for the small sample size) based on QQ plots as well as Shapiro-Wilk test (p>0.05). For “mean distance to WSM vertex”, the sixth subject (Cohort A) was an outlier and was excluded from assessing that feature. A two-tailed paired sample t-test was then used on within-subject means in all cases except for “coverage” which was assessed as an absolute deviation from 1 using a one-tailed paired sample t-test.

##### Effects of scanner, acquisition, resampling

Linear mixed effect modelling was used because the data were non-independent. The model was constructed as follows:

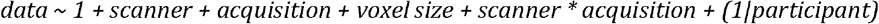

‘Scanner’, ‘acquisition’, ‘voxel size’ were nominal variables representing fixed effects, while ‘participant’ was a nominal variable representing random effects and reflecting the assumption that within-participant data will be correlated. This model produced better estimates for all types of data (confirmed with Bayesian information criterion and likelihood ratio test, the latter based on maximum likelihood estimates) compared against a simpler model without the scanner-acquisition interaction. Interaction with ‘voxel size’ was not considered because upsampling was a post-processing step done by re-gridding and interpolation independent of hardware and scanning protocol. Final estimates were calculated using restricted maximum likelihood. Residuals were inspected for normality (QQ plots and Shapiro-Wilk test), homoscedasticity (by plotting against fitted values), absence of autocorrelation (by plotting in order) to confirm that the model assumptions were met, although a small number of outliers were accepted when assessing for normality. A Box-Cox power transformation on the response variable 3 (“coverage”) was applied to alleviate heteroscedasticity.

##### Consistency of SAF tractograms

Repeatability and between-subject variability were calculated using within- (CV_W_) and between- (CV_B_) subject coefficients of variation, respectively (Laguna et al. (2020)). Reliability of metrics was characterised using single measurement intraclass correlation coefficient for absolute agreement ICC(3,1) with subject effects modelled as random and session effects fixed (McGraw and Wong (1996)). The data were formulated with a linear mixed-effects model (Chen et al. (2018)). For voxel-based (TDI) and surface-based analysis, CV_W_, CV_B_ and ICC were calculated at each voxel/vertex.

### 3.3. Data/code availability statement

- Data: please refer to Koller et al. (2020) for access to the test-retest data, and to Tax et al. (2019) for access to the cross-scanner and cross-protocol diffusion MRI data harmonisation database.
- Code: the MATLAB code for filtering of SAF and interfacing with the surface will be made available upon publication at: https://github.com/dmitrishastin/SAF

## 4. Results

### 4.1. Comparison with voxel-based method

#### Seed distribution

The distribution of seeds that resulted in streamlines is illustrated in relation to the WSM in the top row of Figure 2. This demonstrates that surface seeding produced a smaller number of unique seeds yet appeared to provide a more consistent GMWMI coverage observed in both gyri and sulci. Despite this, a small minority of vertices did not give rise to streamlines.

**Figure 2:**
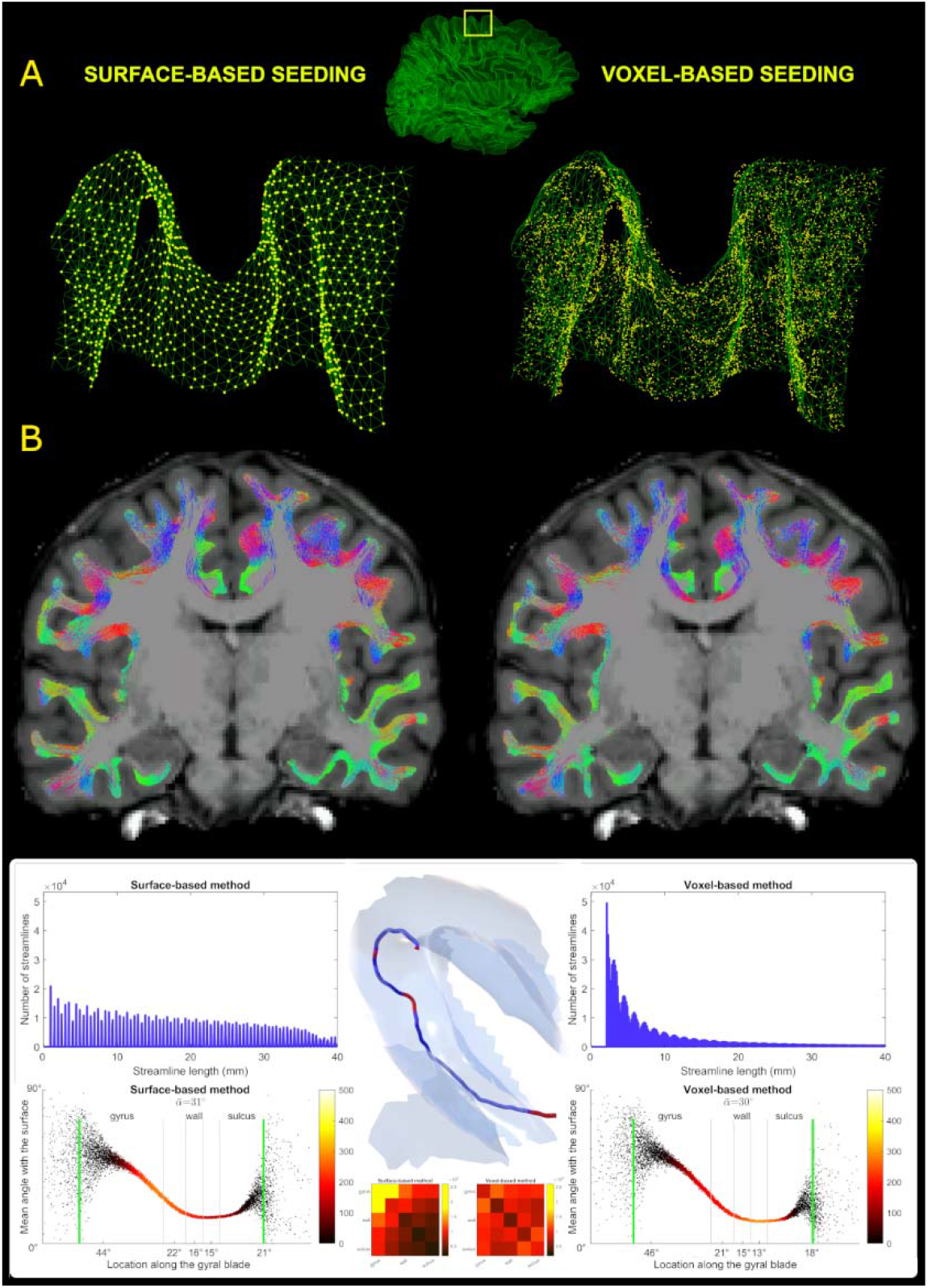
Comparison of surface- and voxel-based strategies. A: Distribution of seeding coordinates (yellow) related to a section of the WSM (green) in the paramedian premotor cortex (region-defining box in the middle). With surface seeding, most (but not all) vertices resulted in streamlines, visually achieving a more spatially uniform distribution. **B:** Final tractograms for the same subject showing similar appearances except for minimal streamline extension in a few areas into the deep grey matter with the surface method. **Bottom, upper row:** Histograms of streamline length (averaged across cohort) show a near-linear decline with the surface method and roughly a power law decline with the voxel method. Illustration in the centre shows a streamline coursing tangential to the grey-white interface (semitransparent). Most sections are subcortical (blue) but the few escapes into the grey matter (red) would have met the conventional termination criteria resulting in a considerably shorter streamline. **Bottom, lower row:** Plots of streamline angle at the cortex along the gyral blade (averaged across cohort) show less acute angle with the surface method. Also see legend for Figure 6. Connectivity matrices in the centre show the number of streamlines connecting the five zones along the gyral blade (averaged across cohort) produced with the two methods. Colour represents streamline counts.

#### Final tractograms

Compared to the voxel-based method (Table 4), the surface approach resulted in longer streamlines (p<0.001) with lengths distributed more evenly across the target range (Figure 2, histograms). Streamlines terminated closer to the surface (p=0.001), approached a larger number of vertices (p<0.001) and these vertices were more evenly distributed between gyri and sulci (p=0.021). In contrast, the voxel-based method offered a far less biased distribution of streamline terminations between gyri and sulci (p<0.001) and had slightly higher mean fractional anisotropy (p<0.001). Additionally, the mean streamline-cortex angle was slightly greater with the surface-based approach (Figure 2, bottom plots). Examining region-specific angles, the same was true in the walls and sulci but the angle in the gyri was greater with voxel-based approach.

**Table 3:**
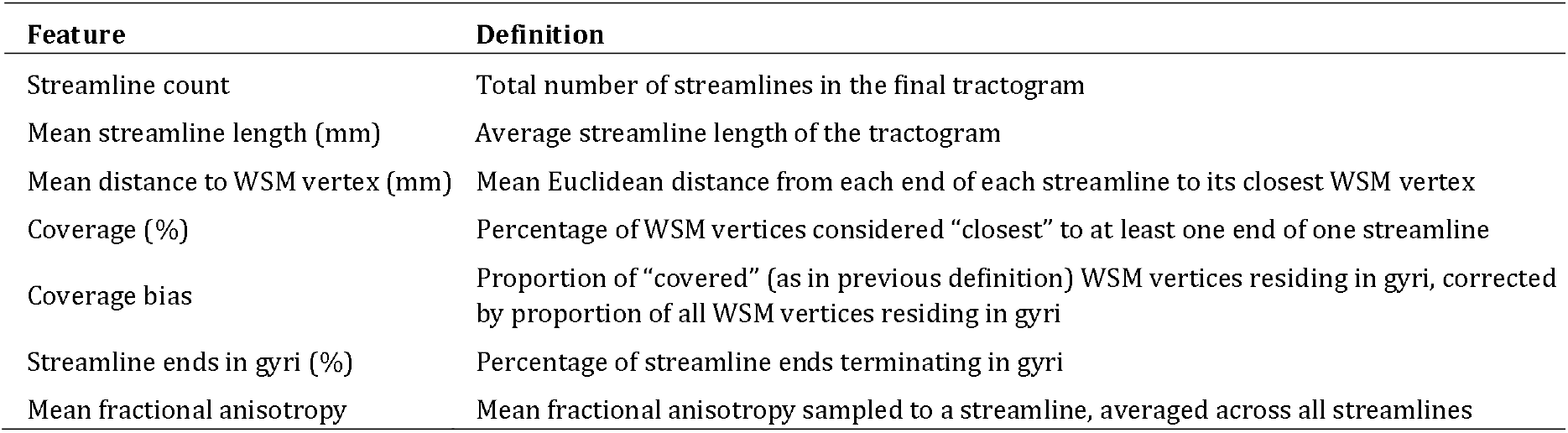
Tractogram features used for evaluation and their definitions.

| Feature | Definition |
| --- | --- |
| Streamline count | Total number of streamlines in the final tractogram |
| Mean streamline length (mm) | Average streamline length of the tractogram |
| Mean distance to WSM vertex (mm) | Mean Euclidean distance from each end of each streamline to its closest WSM vertex |
| Coverage (%) | Percentage of WSM vertices considered “closest” to at least one end of one streamline |
| Coverage bias | Proportion of “covered” (as in previous definition) WSM vertices residing in gyri, corrected by proportion of all WSM vertices residing in gyri |
| Streamline ends in gyri (%) | Percentage of streamline ends terminating in gyri |
| Mean fractional anisotropy | Mean fractional anisotropy sampled to a streamline, averaged across all streamlines |

**Table 4:**
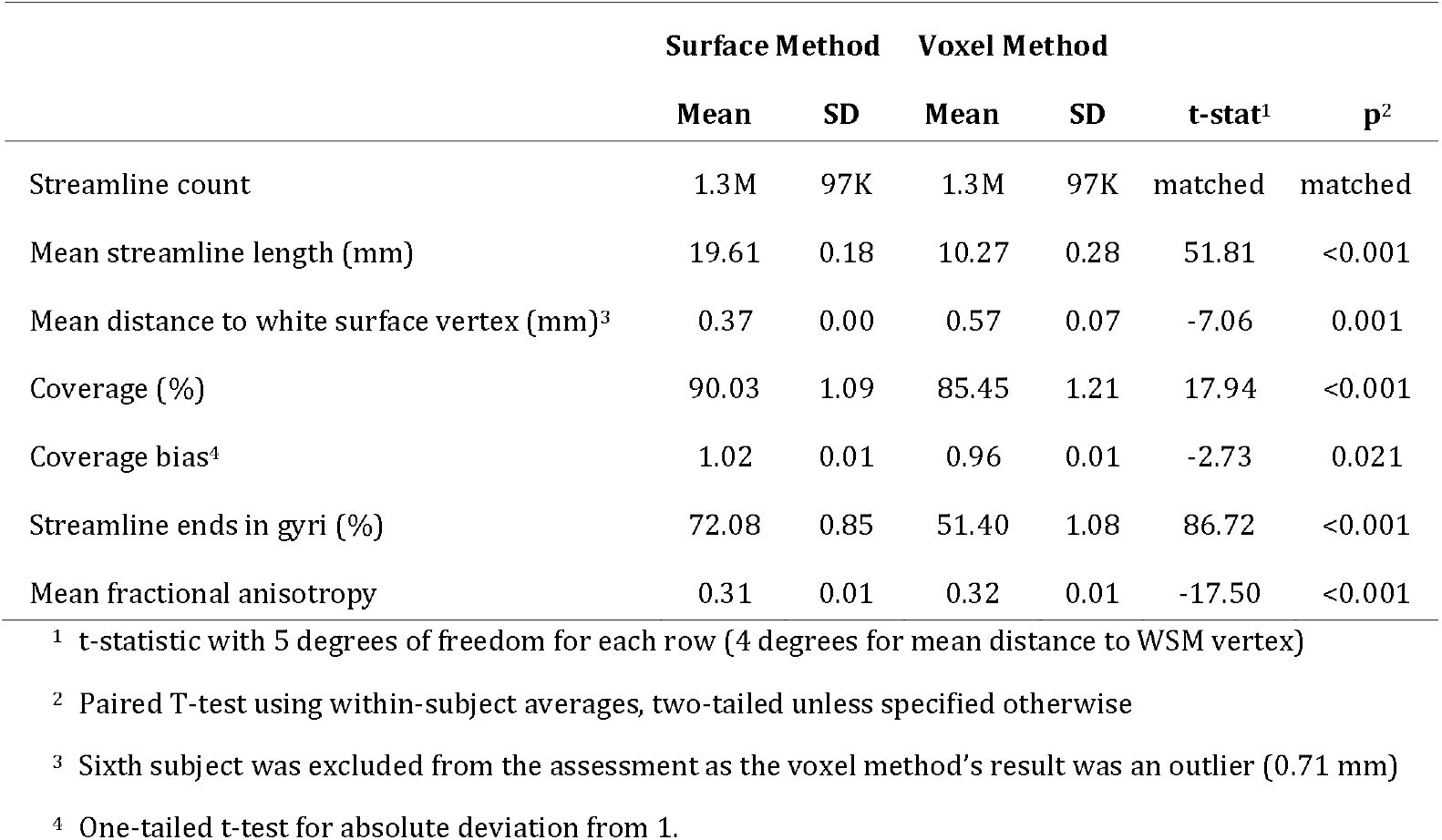
Tractogram differences between surface- and voxel-based methods.

|  | Surface Method |  | Voxel Method |  | t-stat<sup>1</sup> | p<sup>2</sup> |
| --- | --- | --- | --- | --- | --- | --- |
|  | Mean | SD | Mean | SD |  |  |
| Streamline count | 1.3M | 97K | 1.3M | 97K | matched | matched |
| Mean streamline length (mm) | 19.61 | 0.18 | 10.27 | 0.28 | 51.81 | <0.001 |
| Mean distance to white surface vertex (mm)<sup>3</sup> | 0.37 | 0.00 | 0.57 | 0.07 | -7.06 | 0.001 |
| Coverage (%) | 90.03 | 1.09 | 85.45 | 1.21 | 17.94 | <0.001 |
| Coverage bias<sup>4</sup> | 1.02 | 0.01 | 0.96 | 0.01 | -2.73 | 0.021 |
| Streamline ends in gyri (%) | 72.08 | 0.85 | 51.40 | 1.08 | 86.72 | <0.001 |
| Mean fractional anisotropy | 0.31 | 0.01 | 0.32 | 0.01 | -17.50 | <0.001 |

<sup>1</sup> t-statistic with 5 degrees of freedom for each row (4 degrees for mean distance to WSM vertex)
<sup>2</sup> Paired T-test using within-subject averages, two-tailed unless specified otherwise
<sup>3</sup> Sixth subject was excluded from the assessment as the voxel method's result was an outlier (0.71 mm)
<sup>4</sup> One-tailed t-test for absolute deviation from 1.

#### Streamline cropping

It was hypothesised that the difference in histograms of streamline length was driven by differences in termination and/or filtering criteria. The surface approach only truncated the terminal portions of each streamline where they extended into the cortex while everything in between remained unchanged. Filtered this way, it was noted that streamlines would sometimes escape into the cortex in between these terminal portions but promptly return to the white matter after a short distance. Without producing grossly aberrant shapes, this enabled streamlines to contour the grey-white interface for longer. This is illustrated in the 3D model at the bottom of Figure 2 - the intra-cortical segments (in red) are short, near-tangential to the surface and they do not deviate far. On the other hand, the voxel-based method did not allow streamlines to course through the cortex, leading to multiple interrupted, short segments. To confirm this was the key difference, the first data set from each subject’s time series was processed analogously to the main pipeline, except now streamlines were cropped every time the WSM was traversed. Continuing with the illustration in Figure 2, the stricter cropping mechanics broke the streamline into three shorter (blue) streamlines by excluding the red segments. This cropping method produced an average of 2M streamlines with the mean length of 8.76 mm, demonstrating results very comparable to the voxel method. As all processing steps up to the GWG filter were identical, this difference is directly attributable to cropping. Another major similarity with the voxel method was that 51.5% of streamline ends terminated in the gyri, contrasting sharply against the 72.08% seen when cropping at terminations only. Together, these findings strongly suggest that cropping at terminations alone enabled streamlines to “navigate” sulcal fundi more easily with more streamlines reaching the neighbouring gyri therefore translating to increased bias. The connectivity matrices at the bottom of Figure 2 corroborate this by showing that gyri were joined by many more streamlines compared to sulci.

To verify that the streamlines did not veer deeply into the grey matter when only their outer sections were truncated, the white matter mask was dilated by 1 mm^3^ (to include the WSM which was used for cropping). Across all available data sets, on average only 1.47% (range: 0.83-2.15%) of streamlines escaped this dilated mask and only 2.51% (range: 1.40-3.41%) of SAF-containing voxels were found outside, confirming that throughout their length, >95% of streamlines stayed approximately within the confines of the WSM (allowing for the discrepancy between the mesh and the voxel grid).

#### Surface seeding

To study the role of surface seeding specifically, the first data set from each subject’s time series was processed analogously to the main pipeline but the seeding was done from the voxel method’s grey-white interface file using MRtrix *tckgen’s -seed_rejection* flag, meaning the probability of seeding from a given voxel was proportional to the voxel’s value. On average, this resulted in 1M slightly shorter (18.48 mm) streamlines with reduced surface coverage (84.31%) but a marginally decreased gyral bias (68.68%). This was not strictly representative of the voxel method’s seeding mechanic which was difficult to replicate precisely; however, it raises the possibility that surface seeding contributed to the difference in cortical coverage observed in Table 4.

### 4.2. Effects of scanner, acquisition, resampling

The experiments in the following section continue to employ the proposed surface-based seeding and filtering method.

#### Fibre orientation distribution functions

While the shape and direction of most glyphs at the grey-white interface did not differ strongly from a visual perspective, some glyphs along the gyral wall had noticeably more anisotropy and radial orientation with the “state-of-the-art” data, particularly Connectom scanner; the same was even more evident in the cortex (see Figure 3).

**Figure 3:**
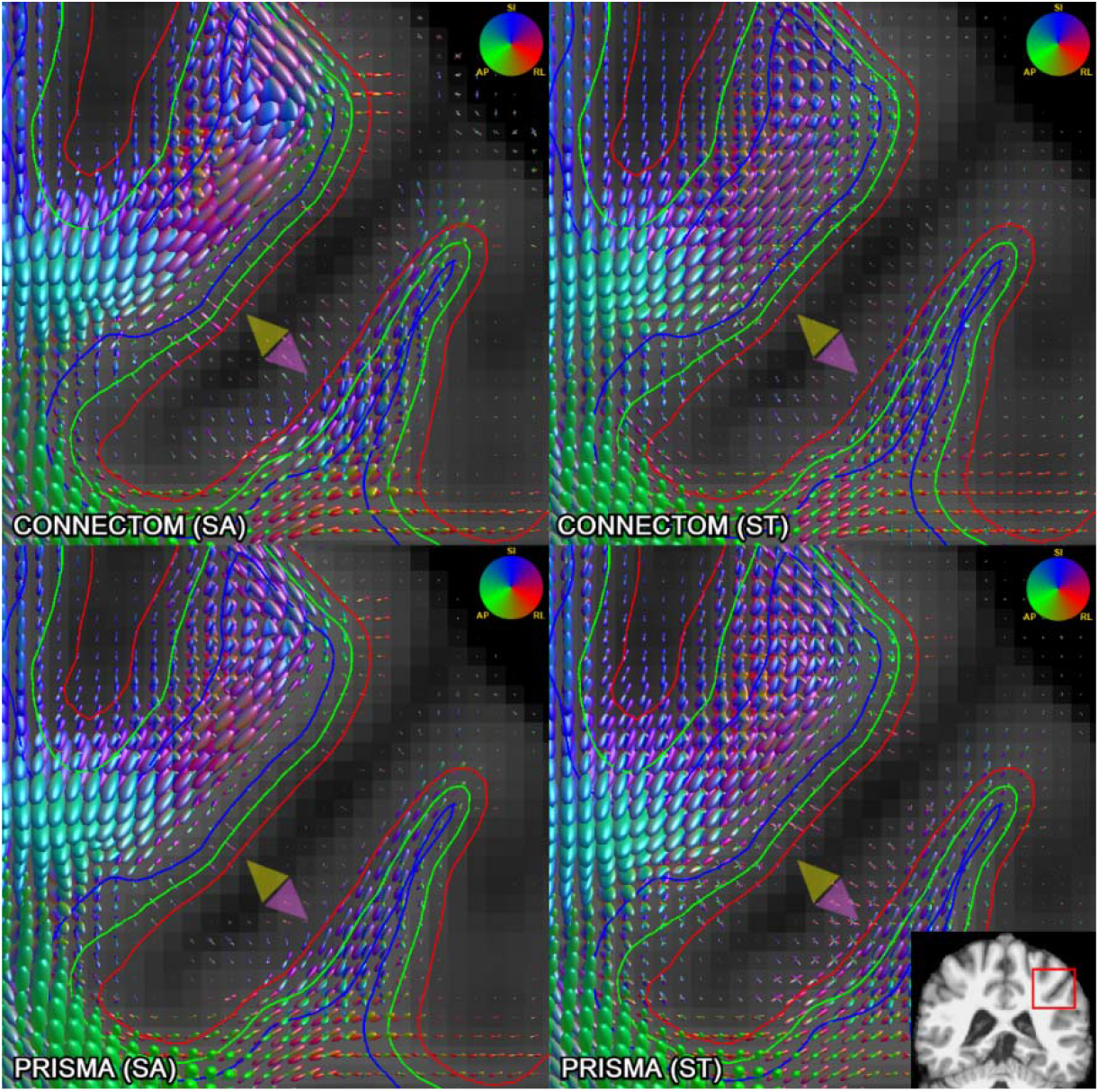
Comparison of fODFs (l_max_=8, scale=3) generated using Connectom (top row) and Prisma (bottom row), “state-of-the-art” (left column) and “standard” (right column) acquisitions for the same subject on a 1 mm isotropic voxel grid, affinely co-registered to the same T1-weighted volume. Inset in the bottom right shows the region compared. Green outline: grey-white interface, red outline: 1 mm superficial, blue outline: 1 mm deep. State-of-the-art acquisitions (particularly from the Connectom scanner) demonstrate increased anisotropy along the gyral wall with glyphs pointing radially (yellow arrowheads) or turning more sharply (purple arrowhead). Colour of the glyphs represents orientation: SI, superior-inferior; AP, anterior-posterior; RL, right-left.

Whole-brain, 1^st^ fODF peak angle plots performed in subject space for all subjects confirm this observation, demonstrating increasingly radial orientation of peaks from deep white matter to cortex; again, “state-of-the-art” sequences showed by far the steepest increase in radiality going outwards, followed by smaller-voxel data and the use of Connectom scanner.

#### Tractogram comparison

There were clear qualitative differences in tractogram appearance depending on the scanner, acquisition, and resampling used (Figure 5). Most notably, for some configurations there existed regions less densely populated by streamlines. The “standard” sequences (lower angular and spatial resolution, lower b-values, lower number of shells) seemed particularly prone to this limitation. Using lower spatial resolution data as well as the Prisma also appeared to contribute to this effect, although to a lesser degree. Further, depending on the scanner and sequence type used, the dominant direction of streamlines appeared to change in the crowns of some gyri.

**Figure 4:**
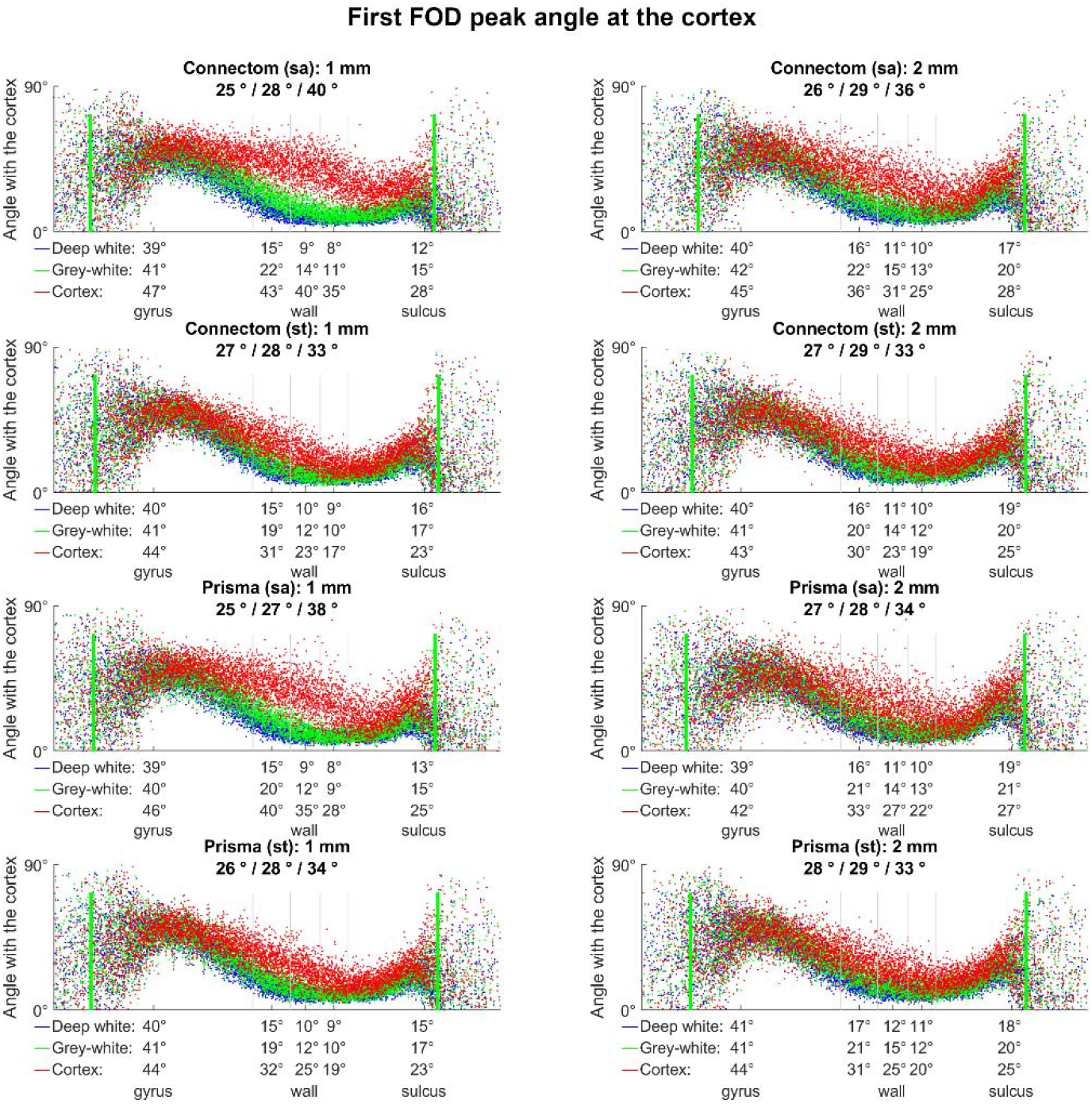
Effect of scanner, acquisition, resampling on the orientation of FODs (calculated for 1^st^ peak only) in relation to the cortex, depending on the location along the gyral blade. Each plot shows average angle distributions across all subjects, sampled at three depths (green: grey-white interface, blue: 1 mm deep, red: 1 mm superficial). Mean angle is shown under each title (deep white/grey-white/cortex, respectively); same is provided on the x-axis for the five equal-area sectors of the gyral blade. Grey vertical lines illustrate boundaries between the five sections along the gyral blade. Green vertical lines represent 5-95% range used for average angle calculation. Connectom scanner, “state-of-the-art” sequences, smaller voxel data all demonstrate steeper peak rotation from near-parallel to near-perpendicular orientation (from deep white to cortex, respectively) particularly in

**Figure 5:**
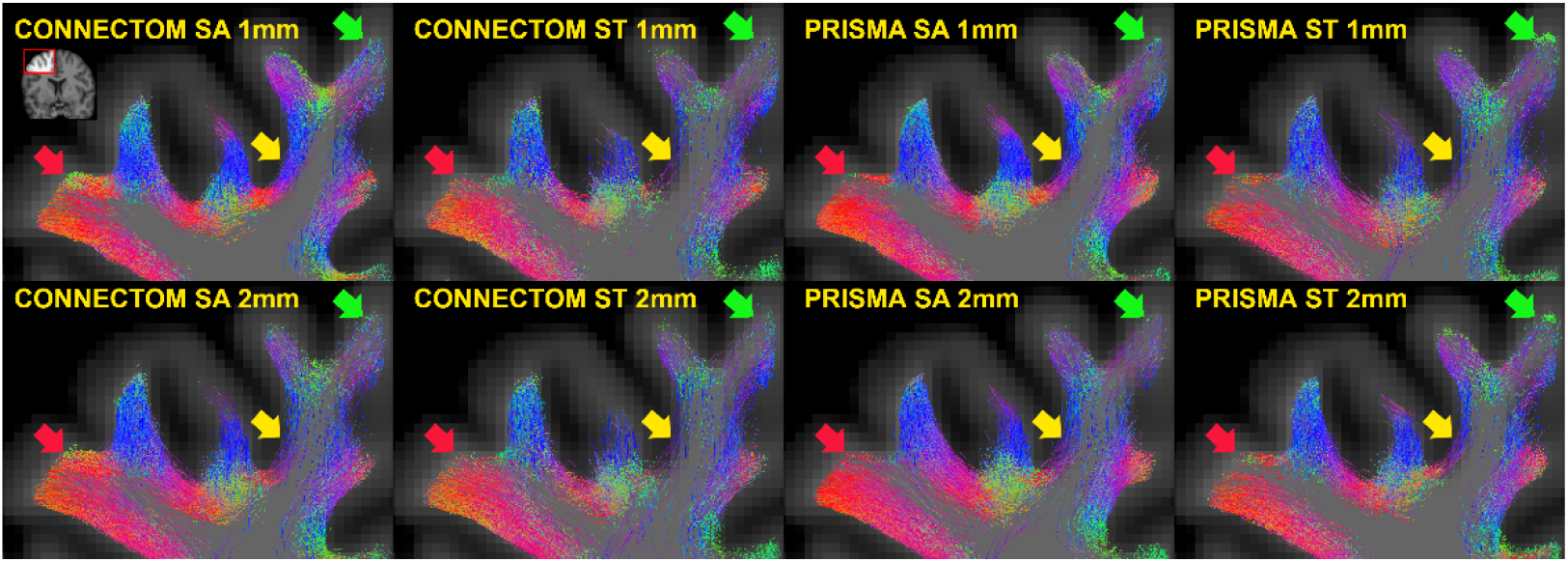
Appearances of SAF tractograms depending on the choice of scanner, acquisition, resampling. A region from the right frontal lobe of the same subject is shown in coronal plane (inset in the left upper corner). Tractogram slices are 1 mm thick, all voxels are isotropic. Differences in streamline density are most noted along a sulcal wall (yellow arrow) and a gyral crown (red arrow). Local variation in the predominant direction of streamlines is also seen (green arrow, red arrow).

The linear mixed model revealed that the choice of scanner, acquisition and resampling all had statistically significant effects on the tractograms (Table 5). Overall, in this cohort the sequence type and data resampling had the most influence on results, while the scanner choice appeared less important. Most notably, the streamline count was profoundly reduced when using “standard” sequences and resampling to larger voxel grids. While there was a statistical difference in mean streamline length with the use of Prisma scanner and larger voxels, this was in the order of 1-2 mm as opposed to the difference with the voxel method (see section 4.1, “Comparison with voxel-based method”). The use of Prisma scanner, “standard” sequences and larger voxels all reduced cortical coverage as well as the number of terminations in the gyri. Finally, mean fractional anisotropy decreased with Prisma and “standard” sequences but increased with larger voxels, possibly as more signal from the deep white matter was included.

**Table 5:** Effect of scanner, acquisition, resampling on the final SAF tractogram estimated using linear mixed effects modelling. Intercept is the mean for Connectom, SA (“state-of-the-art”) acquisition, smaller voxels (reference), and subsequent beta values are estimated fixed effect coefficients (representing deviations from that mean for Prisma, ST (“standard”) acquisition, larger voxels, respectively). The 95% confidence interval is provided in the adjacent columns.

| **s** | **Beta** | **Lower 95%** | **Upper 95%** | **p-value** |
| --- | --- | --- | --- | --- |
| Intercept | 1.25M | 1.19M | 1.30M | <0.001 |
| Scanner (Prisma) | 18K | -32K | 68K | 0.475 |
| Acquisition (ST) | -340K | -390K | -290K | <0.001 |
| Voxel size (2×2×2 mm<sup>3</sup>) | -309K | -344K | -273K | <0.001 |
| Scanner (Prisma) × Acquisition (ST) | 45K | -26K | 115K | 0.213 |
| **Mean streamline length (mm)** |  |  |  |  |
| Intercept | 17.87 | 17.65 | 18.08 | <0.001 |
| Scanner (Prisma) | 1.04 | 0.83 | 1.24 | <0.001 |
| Acquisition (ST) | 0.24 | 0.04 | 0.45 | 0.019 |
| Voxel size (2×2×2 mm<sup>3</sup>) | 2.08 | 1.93 | 2.22 | <0.001 |
| Scanner (Prisma) × Acquisition (ST) | -1.08 | -1.37 | -0.80 | <0.001 |
| **Coverage (%) – power transformed ($\lambda=4.3$)** |  |  |  |  |
| Intercept | -0.08 | -0.09 | -0.07 | <0.001 |
| Scanner (Prisma) | -0.01 | -0.02 | 0.00 | 0.004 |
| Acquisition (ST) | -0.03 | -0.04 | -0.02 | <0.001 |
| Voxel size (2×2×2 mm<sup>3</sup>) | -0.03 | -0.03 | -0.02 | <0.001 |
| Scanner (Prisma) × Acquisition (ST) | 0.03 | 0.02 | 0.04 | <0.001 |
| **Streamline ends in gyri (%)** |  |  |  |  |
| Intercept | 0.73 | 0.73 | 0.74 | <0.001 |
| Scanner (Prisma) | -0.01 | -0.01 | 0.00 | 0.037 |
| Acquisition (ST) | -0.04 | -0.04 | -0.03 | <0.001 |
| Voxel size (2×2×2 mm<sup>3</sup>) | -0.03 | -0.03 | -0.02 | <0.001 |
| Scanner (Prisma) × Acquisition (ST) | 0.02 | 0.01 | 0.03 | <0.001 |
| **Mean fractional anisotropy** |  |  |  |  |
| Intercept | 0.32 | 0.31 | 0.32 | <0.001 |
| Scanner (Prisma) | -0.01 | -0.01 | -0.01 | <0.001 |
| Acquisition (ST) | -0.04 | -0.05 | -0.04 | <0.001 |
| Voxel size (2×2×2 mm<sup>3</sup>) | 0.02 | 0.02 | 0.02 | <0.001 |
| Scanner (Prisma) × Acquisition (ST) | 0.02 | 0.02 | 0.03 | <0.001 |

Contribution of intermodal registration differences to resampling effects arising due to different voxel size was studied qualitatively (Appendix E). A degree of misalignment between 1mm and 2mm data irrespective of scanner or sequence type was obvious; however, the region particularly affected on Figure 5 (yellow arrow) remained well aligned (and vice versa), suggesting that registration imperfections did not dominate these results. Additionally, to assess whether resampling effects were linked to step size during tracking, tractography for a small number of data sets with 2×2×2 mm^3^ voxels was repeated using a halved (0.5 mm) step size; this, however, produced similar results (data not shown).

#### Streamline angle at the cortex

The general shape of the distribution remained similar irrespective of the parameters, with least acute angles seen in the gyri, a near-parallel course along the walls followed by a slight increase in angles near the sulci (Figure 6). For all data sets, the gyral and sulcal angles were more dispersed compared to the walls. While parameter-associated variation of mean angles in most sectors stayed within a degree, in the sulci this increased prominently when “standard” sequences and larger voxels were used, less obviously impacted by scanner choice.

**Figure 6:**
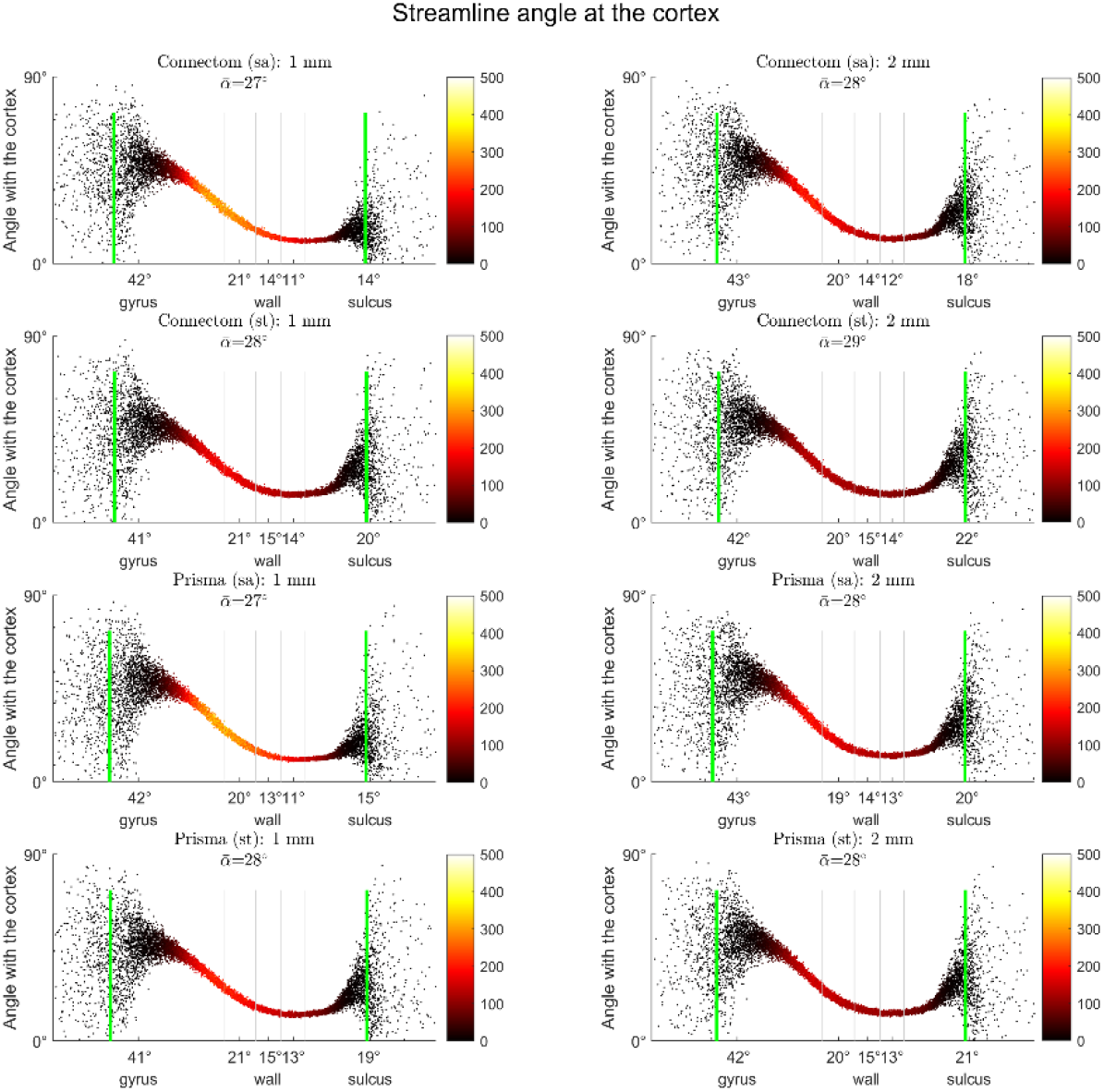
Effect of scanner, acquisition, resampling on the streamline-cortex angle, depending on location along the gyral blade. Each plot shows average angle distributions across all subjects. Colour bars represent streamline counts. Mean angle (not adjusted for streamline count) is shown in each title as 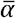; same is provided on the x-axis for the five equal-area sectors of the gyral blade. Grey vertical lines illustrate boundaries between the sectors. Green vertical lines represent 5-95% range used for average angle calculation. Most differences appear in the sulci, with Prisma scanner, “standard” sequence and larger voxels leading to increased mean angles.

#### Distribution of connections along the gyral blade

Similar to Cohort A, connectivity matrices showed that streamlines forming connections from gyri to (same or other) gyri were the most prevalent (Figure 7). This was followed by streamlines connecting gyri to walls and gyri to sulci, with streamlines connecting sulci to sulci being the least common. These differences remained broadly similar irrespective of the parameters used, although their scale increased with Connectom scanner, “state-of-the-art” sequences, and data upsampling. As all data sets started with the same number of unfiltered streamlines and the filtering was performed in the same way, the differences are likely due to how the streamlines propagated at the integration step. Figures 3 and 4 demonstrate that the “state-of-the-art” sequence on the Connectom scanner had more above-threshold fODFs deeper in the cortex of the gyri and walls but not the sulci, with a more anisotropic, radially directed orientation of fODF lobes. This could have resulted in more streamlines reaching the cortex in those segments and thereby surviving the filtering, explaining the differences in final streamline counts and their distribution.

**Figure 7:**
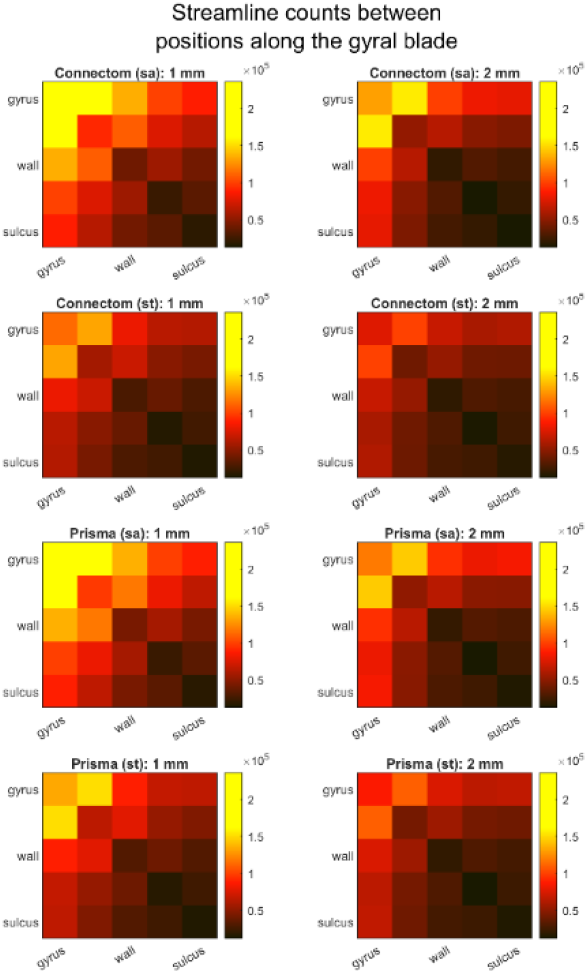
Effect of scanner, acquisition, and voxel size on SAF connectivity between different positions along gyral blades. Cortical surface was divided into five zones with equal areas from sulcal fundi to gyral crowns. Each image shows the absolute number of streamlines connecting different zones across all subjects. Connectivity matrices suggest that Connectom, “state-of-the-art” sequence, small voxel size increased the overall number of connections but particularly so between gyral crowns and the surrounding areas.

### 4.3. Assessment of SAF tractograms

Following the initial tractogram generation, the framework runtime was

2-2.5 hours with parallel processing (12 CPUs) enabled. On average, 26% of the original streamlines survived the filtering. Tractograms in Cohort A appeared to agree with the expected anatomy and no manual pruning was required (Figure 8). For the most part, SAF appeared to course in a bundled fashion and there appeared to be numerous mixing of bundles as highlighted by direction-encoded colour maps (Figures 2, 5, 8). Tractogram features of interest were already presented in section 4.1 (“Comparison with voxel-based method”). The distribution of values on the surface in a single subject is demonstrated seen in Appendix D.

**Figure 8:**
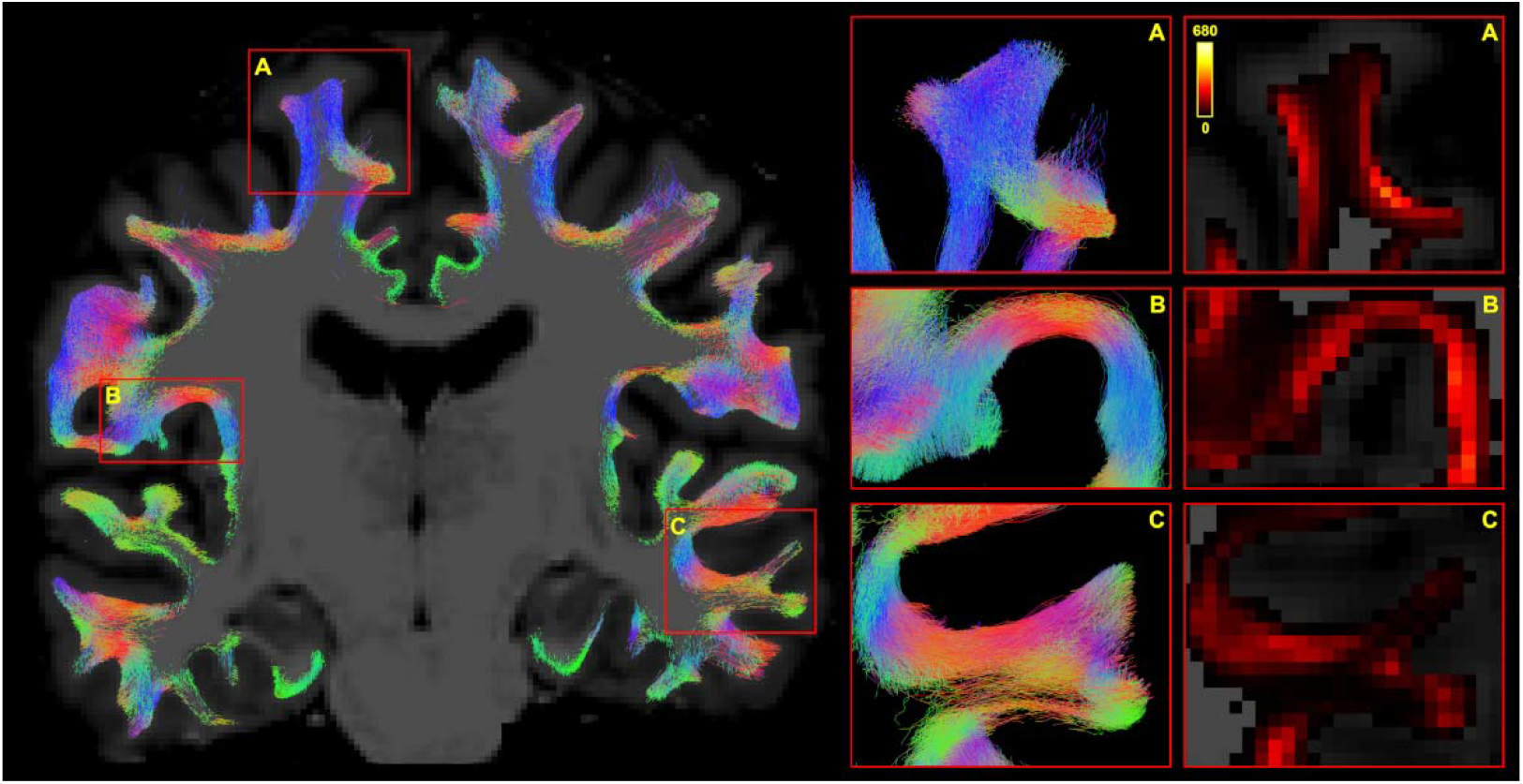
Final appearance of SAF tractogram following the filtering process. **Left:** SAF streamlines overlaid on the T1-weighted volume in dMRI space (coronal view, 1 mm thick slice). **Right, first column:** SAF bundles will often deviate from the orthogonal planes in their course. 5 mm thick slices are provided for a better appreciation of their extent. **Right, second column:** TDI maps of the same regions are provided. Regions on the left are represented in the right-hand columns under matching letters.

Compared to the Connectom/SA/1mm voxel data set from Cohort B, number of streamlines (1.30M vs 1.25M), their mean length (19.61 mm vs 17.87 mm) and mean fractional anisotropy (0.31 vs 0.32) as well as cortical coverage (90.03% vs 90.85%) and the percentage of streamline ends in the gyri (72% vs 73%) were all of the same order (allowing for some minor differences). The plots of streamline-cortex angle distributions (Figures 2 and 6) revealed greater mean angles throughout the surface.

### 4.4. Track density imaging maps

Although TDI maps suggested an overall moderate reliability of the spatial distribution of streamlines (median ICC: 0.754), repeatability was low (median CV_W_: 76.02%) suggesting a lot of variation within subjects (Figure 9). Median CV_B_ was 287.64% (thresholded in the figure), attesting to a very high variation between subjects when comparing TDI maps. The immediate subcortical areas showed the least consistency (typical values for CV_W_: 30-50% reaching 200-600%; typical values for CV_B_: 300-400%), possibly due to cortical folding differences (and the associated registration imperfections) coupled with partial volume effects or less reliable tracking on the grey-white interface.

**Figure 9:**
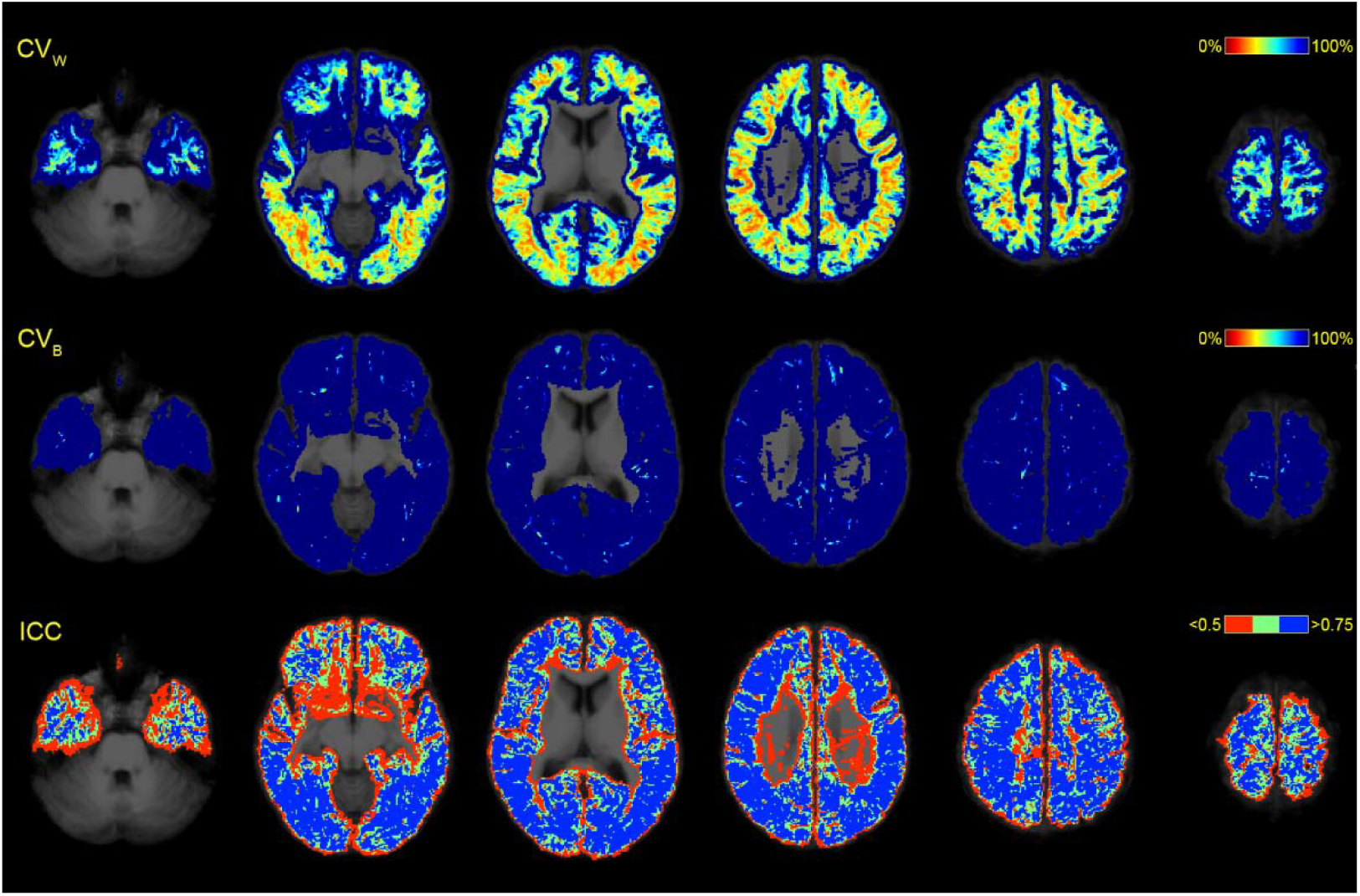
Repeatability of SAF using TDI map comparison in common space. All maps were superimposed on the average T1-weighted volume. CV_W_ and CV_B_ were thresholded at 100%. ICC was thresholded at <0.5 and >0.75. CV_W_, coefficient of variation within subjects. CV_B_, coefficient of variation between subjects. ICC, intraclass correlation coefficient.

To minimise registration-related distortions and cortical folding-driven differences, streamline data were projected on the surface in native space before applying surface registration (Figure 10, also Appendix F). Analysis of termination density demonstrated moderate reliability (median ICC: 0.767) and low repeatability (median CV_W_: 31.55%) together with high between-subject variability (median CV_B_: 106.18%), although this compared favourably to the results seen with TDI analysis. Similar ICC coefficients and improved coefficients of variation were shown for mean length per vertex (median ICC: 0.733, median CV_W_: 13.02%, median CV_B_:40.36%) and mean FA per vertex (median ICC: 0.687, median CV_W_: 8.05%, median CV_B_: 21.70%).

**Figure 10:**
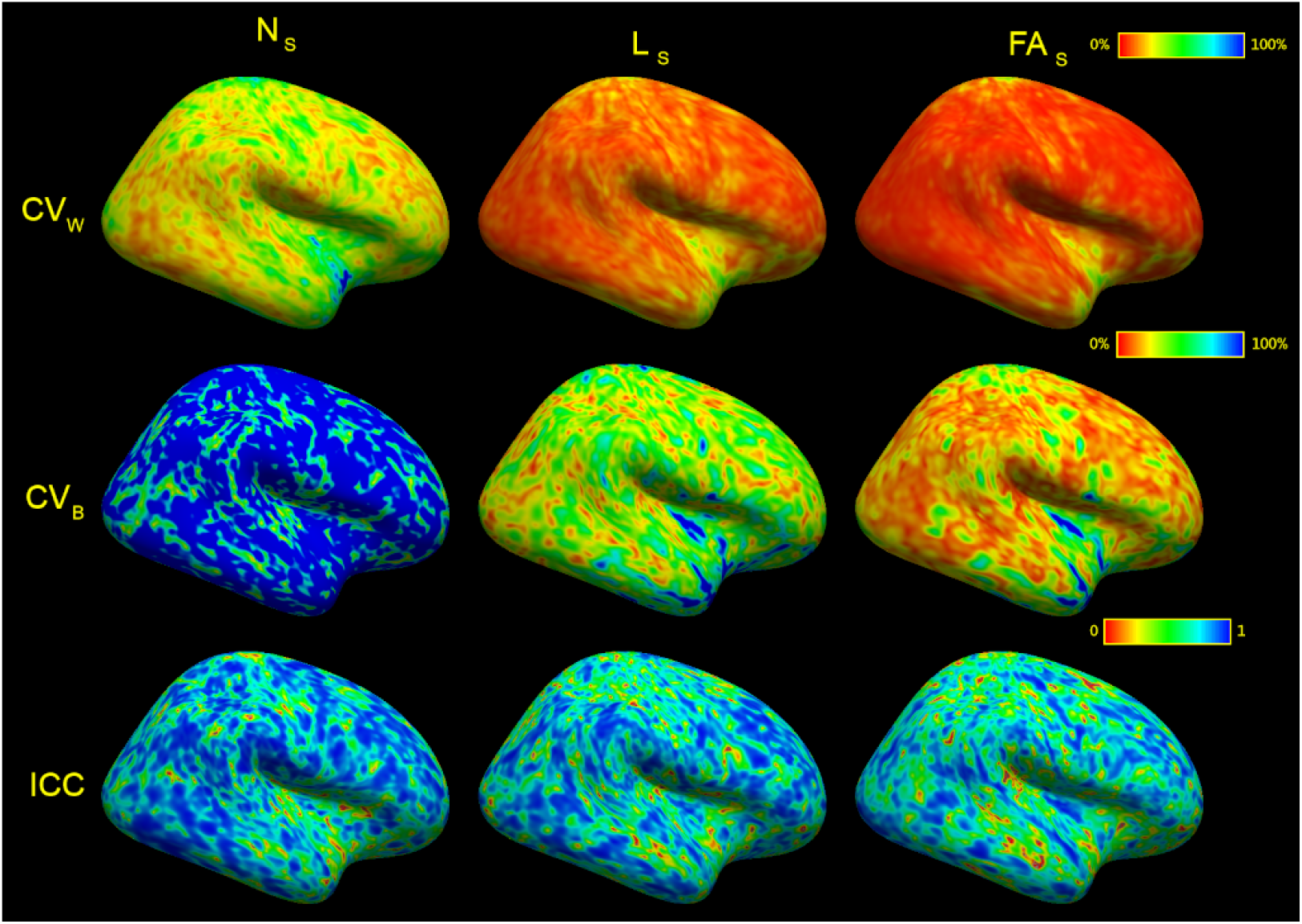
Surface-based analysis demonstrated on the lateral cortex of the right hemisphere. Ns, termination density (number of streamlines/vertex). Ls, mean streamline length/vertex. FA_S_, mean streamline fractional anisotropy/vertex. CV_W_, coefficient of variation within subjects. CV_B_, coefficient of variation between subjects. ICC, intraclass correlation coefficient. Values at each vertex were recorded in subject space, then transformed into average subject space before running analyses. CV_W_ and CV_B_ were

## 5. Discussion

We have presented a focused approach to short association fibre analysis by marrying tractography with mesh representation of the cortex motivated by the close association of SAF with the latter. We were specifically interested in studying the shorter pathways (consistent with the definition in Schüz and Braitenberg (2002) as these pathways are particularly sensitive to inter-individual cortical folding variations (Bajada et al. (2019)) and harder to study using the more established approaches (Román et al. (2017); Zhang et al. (2018); Guevara et al. (2017); Van Essen et al. (2014)). We believe that this study adds a number of useful contributions to the literature.

### 5.1. Comparison with voxel-based method

First, we compared surface- and voxel-based approaches in the context of SAF tractography. We demonstrated that the voxel method produced preferentially short streamlines (mean: 10.27±0.28 mm, median: 6.29±0.27 mm) with a uniform cortical coverage (evidenced by the connectivity matrices and minimal gyral bias). Conversely, the surface method was more inclusive of longer streamlines (mean: 19.61±0.18 mm, median: 19.69±0.36 mm) with predilection for connecting gyri consistent with the common definition of U-fibres (Schüz and Braitenberg (2002)). This was due to the particular filtering strategy we adopted; interestingly, contrary to our initial hypothesis, the main difference did not come from using a surface-based approach specifically but rather from choosing to remove only the terminal intracortical streamline segments, meaning that incursions into the cortex along the streamline course (which we demonstrated to be minimal) were accepted. The effect may potentially be sought after in the context of connectomics, clustering or shape analyses where the very short streamline length may decrease sensitivity to regional changes. On the other hand, along-streamline microstructural sampling will be more confounded by partial volume effects. We explored this in the context of mean fractional anisotropy; and with our small sample (n=6), the difference was marginal (surface mean: 0.31±0.01, voxel mean: 0.32±0.01) but statistically significant (p<0.001). We did not perform comparisons with alternative voxel-based techniques but expect that only truncating the terminal intracortical segments would produce results akin to those of our surface-based method.

Compared to filtering, the choice of seeding strategy, while seemingly less impactful, still merited consideration. In our assessments, surface-based seeding increased the coverage of the grey-white interface (from 84.31% to 90.03%) and resulted in more streamlines (from 1.0M to 1.3M); however, we were not able to replicate the voxel-based seeding strategy precisely. We did not explore the effects of placing more seeds on the surface (for surface-based seeding) or increasing the seed mask resolution (for voxel-based seeding). We would expect the former to further increase cortical coverage (e.g., FreeSurfer WSM is particularly anisotropic around the gyral crowns which was partially addressed with our remeshing strategy and could be further improved using uniform surface sampling (Bowers et al. (2010)), and the latter to decrease the number of streamlines and make them shorter (as fewer seeds will be placed in the subcortical white matter negating the effects described in Reveley et al. (2015)).

### 5.2. Effects of scanner, acquisition, resampling

Second, we investigated the impact of scanner, sequence, dMRI data interpolation on SAF tractograms produced with the surface method. Overall, the Connectom scanner, “state-of-the-art” sequence (more shells, stronger diffusion weighting, higher angular and spatial resolution) and upsampled data had the most satisfactory results. Considered in isolation, sequence type seemed to play the biggest role as the “state-of-the-art” data had more streamlines and greater cortical coverage. This is consistent with previous reports using higher angular resolution (Vos et al. (2016)) and was likely accentuated by using probabilistic tractography (Grisot et al (2021)). Similar changes were observed with upsampling, in keeping with the literature (Dyrby et al. (2014)). The choice of scanner (Connectom with MGA=300 mT/m vs Prisma vs MGA=80 mT/m) impacted the outcomes the least with tractograms demonstrating visually comparable results. This latter finding suggests that the approach may be suitable for good-quality data from clinical scanners, making it more widely accessible.

It remains uncertain whether sacrificing angular in favour of greater spatial resolution would be worthwhile for imaging this particular population of fibres. Previous work has suggested the ideal voxel size of under 0.85 mm isotropic to remain sensitive to the shorter subpopulations of SAF based on the smallest U-fibre turning radius of 0.95 mm (Song et al. (2014); Movahedian Attar et al. (2020)). Considering the angular error of 15-25° in the superficial white matter at 0.25-1 mm resolution across a broad range of q-space sampling schemes and orientation reconstruction methods, the voxels would probably have to be even smaller (Jones et al. (2020)). On the other hand, higher b-values and high angular resolution increase sensitivity to the intra-axonal component of the white matter and allow a better resolution of crossing fibre orientations, respectively (Novikov et al. (2019); Vos et al. (2016); Jeurissen et al. (2014)). Despite their lower spatial resolution, the tractograms of Cohort A appeared to match, if not surpass in quality, those of Cohort B. Nevertheless, fODF-based tractography algorithms may not perform as well in regions with branching, turning, fanning fibres even when multiple fODF peaks are detected (Grisot et al. (2021)). More focused research into this question would be of benefit to mapping SAF in the future. It would also be important to establish whether acquisition schemes and orientation reconstruction approaches that maximise the accuracy of SAF tractography would degrade the fidelity of long tract delineation, and vice versa.

### 5.3. Gyral bias

Third, a special consideration was given to investigating gyral bias in the context of SAF tractography. This was conducted by inspecting tractograms, comparing distributions of streamline terminations and streamline-associated vertices in gyri and sulci, and evaluating streamline connections between sections along the gyral blade. Additionally, we looked at the angle formed by first streamline segments and the overlying cortical mesh depending on the position along the gyral blade; and, in Cohort B, also the angle formed by first fODF peaks (at different depths) and the overlying cortical mesh.

Our filtering strategy produced longer streamlines that largely circumvented sulcal fundi and connected gyri (72.1% terminations in the gyri). Enforcing strict streamline cropping for every WSM intersection produced a termination distribution that was strikingly similar to that of the voxel-based method (51.5% of streamline ends in the gyri). This is significant as we demonstrate that, aside from the underlying anatomy, orientation reconstruction accuracy and tractography integration algorithm, gyral bias is strongly dependent on filtering/termination criteria. Histology suggests that fewer axons enter the cortex around the sulcal fundi; this skew, albeit exaggerated, is also seen with tractography (Schilling et al. (2018)). On the other hand, it is also well known that streamline density is not representative of the underlying axonal density (Schilling et al. (2018)) or even the underlying fODF integrals (Smith et al. (2015)). Depending on the intended application, distributing weights to individual streamlines based on the diffusion signal (Smith et al. (2015); Daducci et al. (2015)) may provide a more biologically meaningful measure of connectivity although this remains to be explored in the context of SAF. For this to be effective, axonal populations need to be represented with at least some streamlines. Tractograms occupied similar anatomical spaces with surface and voxel methods; however, certain areas (particularly near sulcal walls) became underrepresented when the “standard” sequences (as well as, to some degree, larger voxel size and lower MGA scanner) were used. Quantitively, the surface method (p<0.001), “state-of-the-art” sequences (p<0.001), data upsampling (p<0.001) and Connectom scanner (p=0.004) increased cortical coverage.

A secondary observation was that irrespective of acquisition parameters, resampling strategy or tractography approach used, the mean streamline-cortex angle varied only by a few degrees. On this finer scale, “standard” sequences and larger voxels promoted more “perpendicularity” with the cortex in the sulci. This goes against the existing histological evidence whereby axons enter the sulcal cortex at near-tangential angles (Van Essen et al. (2014), Schilling et al. (2018)); these parameters were therefore unwanted. Furthermore, when comparing the orientation of fODF peaks at different depths with streamline orientation, it is clear that the mean streamline orientation at the grey-white interface (in all sections along the gyral blade) largely followed that of the 1^st^ fODF peak at the same depth. The sharper rotation of peaks observed in the cortex of the gyral walls with “state-of-the-art” sequences (as well as Connectom scanner and upsampled data) had no influence on streamline orientation (0.5 mm step size). More generally, while the proportional relationship between different sectors remained consistent with the published histology (Schilling et al. (2018)), the range of absolute values was much lower (41°-46° for crowns, 13°-16° for walls with tractography, 68±15° and 40±17° with histology, respectively). This discrepancy is within the angular error in the superficial white matter shown by Jones et al. (2020) which is also an order greater than its variation arising from different q-space sampling schemes, orientation reconstruction methods and spatial resolution.

### 5.4. Surface representation

Fourth, our approach to the representation of SAF-related measures on the surface merits a mention. Projection of streamline-related data on the surface can be achieved by searching for all streamlines within a sphere around a WSM vertex (Padula et al. (2017); Bajada et al. (2019)). This approach lacks specificity and in regions where non-continuous parts of the cortical mantle lie in proximity with each other (e.g., opposite banks of a narrow gyrus), erroneous inclusion of streamlines may occur. Alternatively, streamline density (Li et al. (2010); Nie et al. (2011)) and orientation termination (Chen et al. (2012)) around a surface vertex have been quantified as the number/orientation of streamlines penetrating the adjacent faces normalised by the combined surface area of the faces. Our approach similarly relies on mesh intersection; however, by lowering fODF threshold (in the context of increased diffusion weighting and higher angular resolution) we enable many more streamlines to reach the cortex. Alternatively, in tractograms where streamlines do not fully approach the mesh, the allocation can be done using the shortest Euclidean distance; we applied this for the voxel-based method by pre-clustering WSM vertices as described in section 3.1.2. Such “projection” on the surface is particularly suited in the context of SAF due to their short length and local course, ensuring each vertex is reasonably representative of its environment. It takes advantage of surface registration and may be used for per-vertex or cluster-based statistical comparison methods, circumventing the use of cortical parcellation if desired and therefore avoiding the associated issues of lower sensitivity within and artificial boundaries between cortical regions.

### 5.5. Consistency of SAF tractograms

Our analysis is complemented by the evaluation of whole-brain SAF tractograms for repeatability, reliability, and between-subject variability. This demonstrated varying results depending on the approach taken. As expected, track density imaging maps demonstrated large variability in the spatial distribution of SAF between individuals but also within individuals; in the attempt to minimise the role of registration imperfections and partial volume effects, alternative measures such as regional density and mean streamline length of SAF were compared by projecting them on the surface resulting in improved repeatability and reduced between-subject variability but similarly moderate reliability. This urges caution when performing analyses that depend on streamline numbers (evidenced by TDI maps and termination density analysis).

Consistency of whole-brain SAF representation with streamlines tractography has previously been addressed. Zhang et al. (2010) emphasised large spatial variability and difficulty in manual region-of-interest segmentation of these tracts advocating for an automated approach. Zhang et al. (2014) demonstrated more short-range than middle-range streamlines with HARDI and DSI data, consistent with our results, and the reverse for DTI data. They proposed the inability of the latter to detect crossing fibres and therefore more false negatives as the likely mechanism. Guevara et al. (2017) used parcellation and shape- and distance-based clustering and found low-to-moderate variability in streamline counts. Román et al. (2017) refined this method by using non-linear registration, allowing detection of within-node connections, clustering larger streamlines and using a bagging strategy, and reported moderate-to-high repeatability of individual bundles. Zhang et al. (2018) generated an atlas of white matter pathways without the use of a cortical parcellation, and subsequently applied it to a number of additional data sets with variable acquisition methods that spanned different age ranges and included clinical cohorts. The authors showed high between-subject variation and moderate overlap between subject and atlas clusters. The atlases (Guevara et al. (2017); Román et al. (2017); Zhang et al. (2018)) were later compared in MNI space for bundle similarity, showing good overlap (Guevara et al. (2020)). The same paper compared the impact of different tractography algorithms on consistency of clustering, showing that probabilistic tracking was able to reconstruct all bundles in 100% of cases with a greater spatial coverage but lower repeatability. All these studies, however, placed focus on longer cortico-cortical connections (starting from 20-35 mm) exclusively, highlighting poor performance with shorter streamlines (Román et al. (2017)). It must also be pointed out that all of the above-mentioned work performed shape-based clustering. Our approach, on the other hand, is agnostic to streamline shape which can be an advantage (being more inclusive) or a disadvantage (being more influenced by false positive streamlines). Shape-based exclusion of noisy streamlines can be implemented (Drakesmith et al. (2019); Parker et al. (2016)); however, the limited availability of validation data for SAF on the whole-brain scale makes identification of such streamlines difficult. At the same time, the data offered here can be considered as a baseline for future methodological developments. Development of better scanning hardware, orientation reconstruction and tractography/optimisation algorithms will also facilitate greater fidelity of the tractograms. The use of multimodal surface registration algorithms (Robinson et al. (2014)) may further improve surface-based consistency.

While not studying SAF on a whole-brain scale, the study by Movahedian Attar et al. (2020) is of special interest as, in common with our approach, it relied on the length definition of Schüz and Braitenberg (2002) and used dMRI data acquired using ultra-high gradient Connectom scanner (choosing higher spatial resolution over stronger diffusion weighting). Using relative streamline counts, the study evaluated connectivity within the occipital cortex. The authors divided the bundles into “retinotopic” (reflecting known anatomy) and “non-retinotopic” (false positive by design) based on fMRI-derived regions-of-interest. They demonstrated high consistency of “retinotopic” bundles (ICC 0.88±0.70, CV 0.23±0.23) but also moderate consistency of “non-retinotopic” bundles (ICC 0.69±0.35, CV 0.25±0.14) which suggests that measures of consistency for SAF are strongly confounded by false positives and again underlines the need for better validation methods on a whole-brain level.

### 5.6. Limitations

Some limitations of our work should be highlighted.

First, despite performing detailed analyses of tractograms our study does not provide histological validation. We referenced existing work (Van Essen et al. (2014), Reveley et al. (2015); Schilling et al. (2018); Yoshino et al. (2020)) to support our results; however, the lack of detailed, whole-brain ground truth makes such correlation incomplete, and this applies to all SAF tractography work to date. It is our hope that this gap will be filled as the interest in mapping the subcortical white matter increases. Understanding the dominant fibre configurations (crossing, bending, fanning, turning) in subcortical white matter will inform the choice of q-space sampling, orientation representation and tractography algorithms (Grisot et al. (2021)); while data on regional variability in length, shape, density of SAF will provide a benchmark for methodological innovation.

Second, the two cohorts used for our analyses had small numbers. The cohorts were chosen because collectively they included unique data from the Connectom scanner allowing the impact of using ultra-high gradient strengths to be assessed and offered multiple timepoints for repeatability analyses. While the results need to be treated with caution, these data allowed to understand parameters that have an important role in SAF tractography, appreciate the limitations of the proposed framework, and identify areas for further development. Our results suggest that, in the context of our framework, SAF reconstructions from good quality data collected on a clinical scanner are comparable to those from data collected on a Connectom scanner. This makes it possible to assess the effects described here in larger cohorts; however, further testing is outside the scope of this paper.

Third, the use of cortical surface meshes is central to our framework and as such it remains sensitive to registration quality between T1-weighted and dMRI data. Data sets containing distortions (such as those arising from susceptibility differences) or unusual anatomy (e.g., tumours) are more likely to suffer from misalignment between the surfaces reconstructed from T1-weighted images and the white matter signal on dMRI. This makes visual inspection on an individual basis crucial. dMRI-based surface extraction could offer an alternative solution (Liu et al. (2007); Li et al. (2010); Shastin et al. (2020)) if performed at sufficiently high resolution.

Last, the filtering pipeline took about 2-2.5 hours to run using parallel CPUs, following the initial tractogram generation. Based on our experiments we judged that such approach is suitable for SAF tractography as exact interaction between streamlines and cortical structures is important. Further developments to the framework could include GPU-based execution and the use of geodesic rather Euclidean distance for cortical coordinate clustering (Lopez-Lopez et al. (2020)).

## 6. Conclusions

We consider our work to be a first-time application of a dedicated whole-brain, surface-based SAF tractography approach aimed at refining the study of this white matter sub-population. Surface-based seeding and filtering algorithms coupled with a streamline length criterion (≤30-40 mm) ensured larger cortical coverage (90%) and promoted gyrus-gyrus connections (72% streamline ends in the gyri) consistent with a common definition of short association fibres. Despite some advantage of using the ultra-high maximum gradient amplitude scanner, the “state-of-the-art” acquisition sequences comprising higher angular resolution and stronger diffusion weighting appeared to achieve comparable results with a clinical scanner making the framework more broadly applicable. “Standard” sequences, however, appeared to underrepresent some subcortical white matter. There also appeared to be an advantage in upsampling the data. At the same time, our voxel-and surface-based evaluations of streamline density showed moderate reliability, low repeatability, and high between-subject variability, urging caution with streamline count-based analyses. Some limitations such as the discrepancy between streamline and axonal trajectories at the grey-white interface continue to present challenge and must be taken into consideration. Overall, the presented framework could be used as a vehicle for investigating SAF in health as well as in clinical cohorts while also offering a platform for future methodological experimentation, while the data presented here can be used for benchmarking.

Declarations of interest: none

## Author contributions

Dmitri Shastin: conceptualization, methodology, software, validation, formal analysis, writing - original draft, writing - review & editing, visualization, funding acquisition. Sila Genc: methodology, software, writing original draft. Greg D. Parker: conceptualization, software. Kristin Koller: resources, data curation. Chantal M.W. Tax: resources, data curation, methodology, writing - review & editing. John Evans: resources, methodology. Khalid Hamandi: writing - review & editing, supervision. William P. Gray: writing - review & editing, conceptualization, supervision. Derek K. Jones: conceptualization, writing - original draft, writing - review & editing, supervision. Maxime Chamberland: conceptualization, writing - original draft, writing - review & editing, visualization, supervision.

## Acknowledgements

This research was funded in whole, or in part, by a Wellcome Trust Investigator Award (096646/Z/11/Z), a Wellcome Trust Strategic Award (104943/Z/14/Z), a Wellcome Trust-funded GW4-CAT fellowship (215944/Z/19/Z), the Engineering and Physical Sciences Research Council (EP/M029778/1), the Dutch Research Council (17331), a Radboud Excellence Initiative Fellowship, and the Brain Repair and Intracranial Neurotherapeutics (BRAIN) Unit funded by Health and Care Research Wales. For the purpose of open access, the author has applied a CC BY public copyright licence to any Author Accepted Manuscript version arising from this submission.

DS would like to thank Prof. Peter H. Morgan at Cardiff Business School for discussions about statistics.

