## Appendix A for "Surface-based tracking for short association fibre tractography"

### Appendix A. Acceptance/rejection of streamlines by GG filter

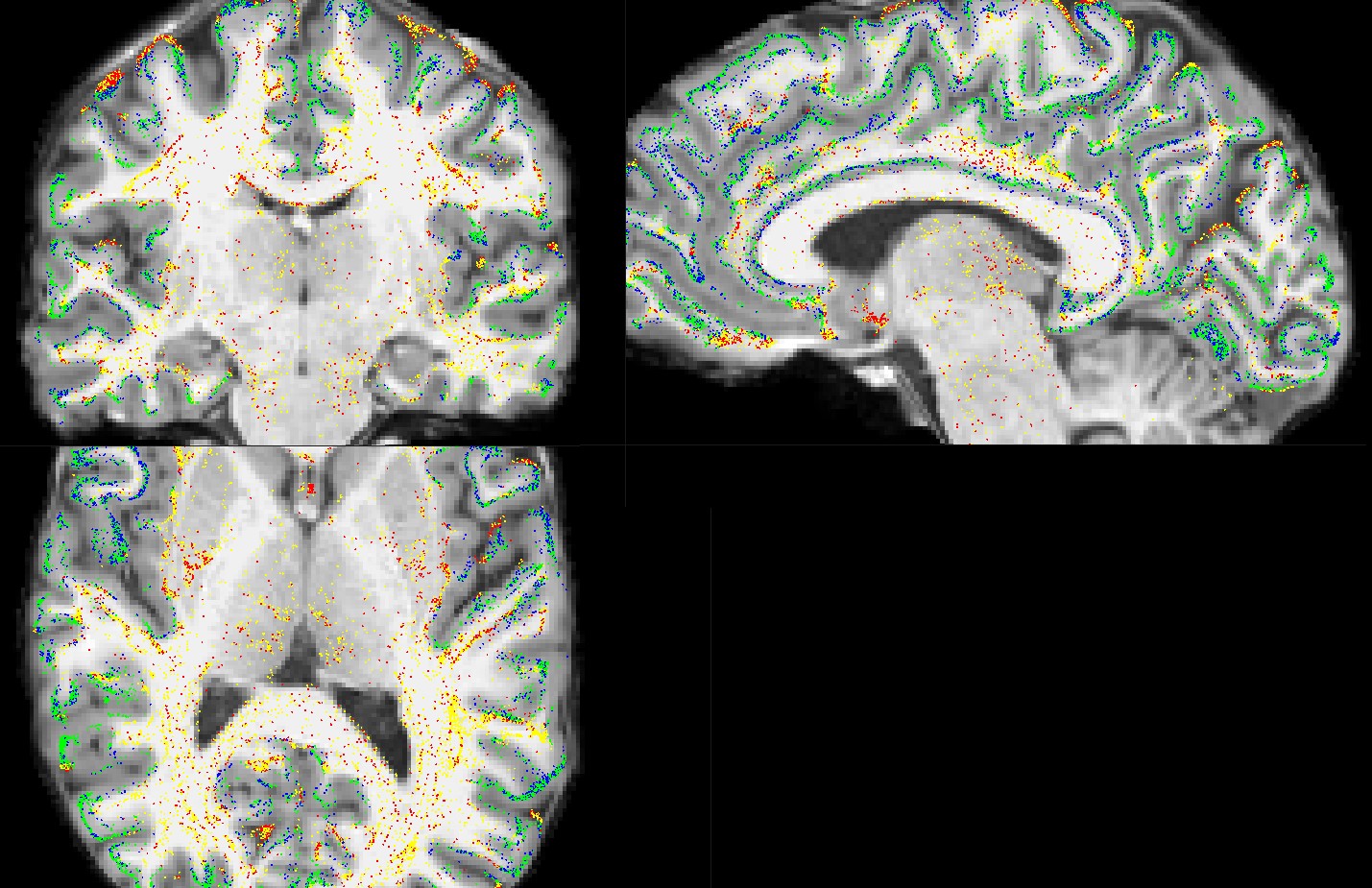

Figure A.1: Both ends of each streamline are demonstrated as points overlaid on T1 image (in dMRI space) in three orthogonal planes. Ends of accepted streamlines are shown in green; all other colours represent rejected streamlines: those with both ends rejected (red) and those with one end rejected (yellow - rejected end; blue - other end). All ends classed by the filter as residing within the cortex (green and blue points) are seen in expected areas; the vast majority of ends rejected by the filter (red and yellow points) are in the white matter, subcortical grey matter, cerebellum or CSF spaces. It is possible that a small proportion of rejections was due to the imperfections of T1-DWI registration. In this early demonstration, 1M seeds were used with no upper limit to streamline length.
