## Appendix B for "Surface-based tracking for short association fibre tractography"

### Appendix B. Description of GG filtering

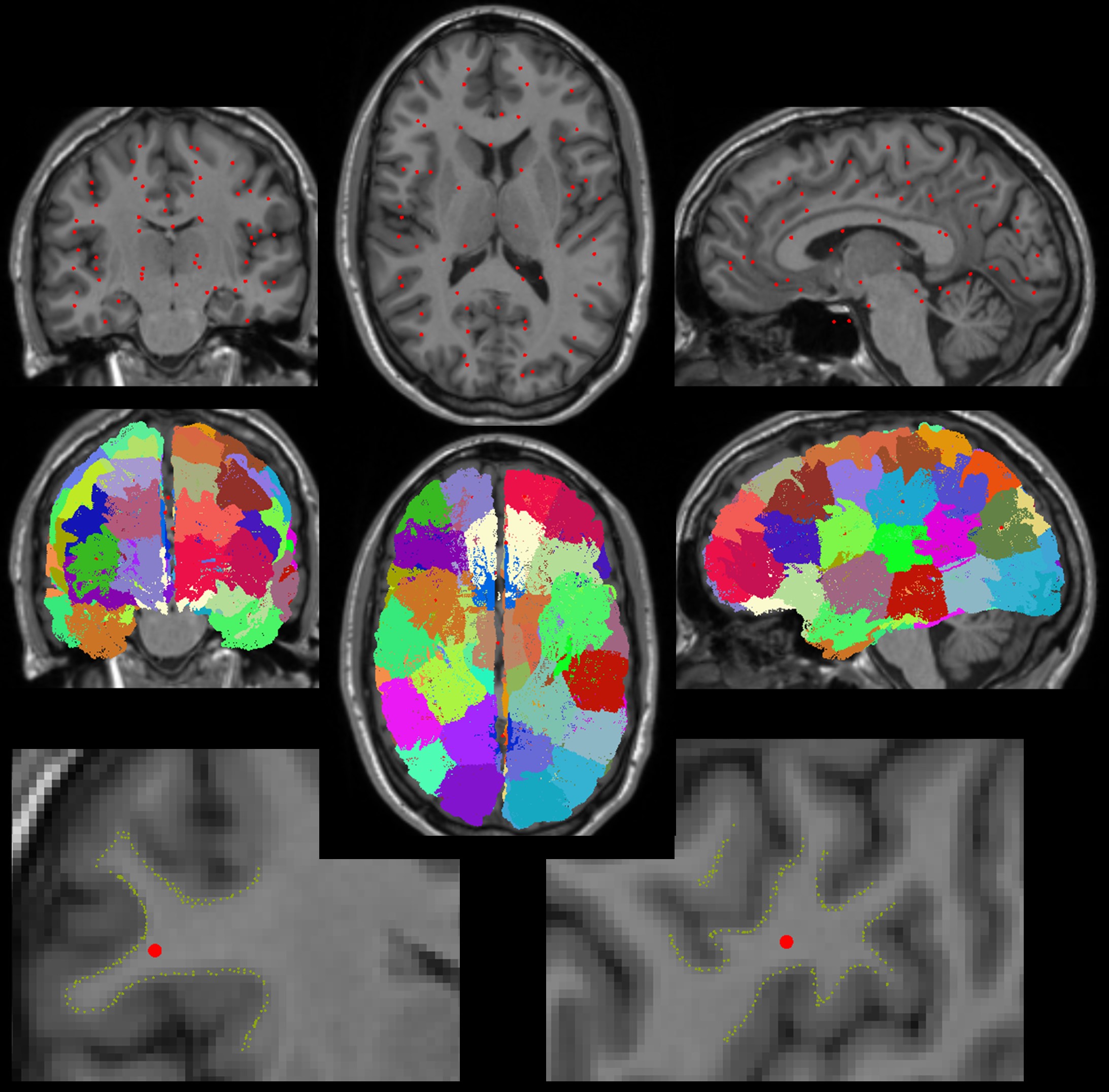

Figure B.1: K-means clustering of vertices (60 clusters were used for illustration purposes). Top row: cluster centroids (red points) projected onto three orthogonal slices of the corresponding T1 volume. Middle row: all vertices colour-coded based on their cluster. While some clusters span both hemispheres (e.g., pale yellow), this does not matter for GG filtering. Bottom row: spatial relationship between a cluster centroid and the vertices belonging to the same cluster.

During GG filtering, Euclidean distances between the coordinates of each end of all streamlines S and the coordinates of all vertices V are calculated, producing a distance matrix with D elements:

$D=2\times S\times V$	(C.1)

The minimal Euclidean is then found for each end of each streamline, corresponding to the vertex closest to that streamline end. To reduce D, the vertices were first clustered into k groups based on their spatial proximity:

Euclidean distances were calculated to each cluster’s centroid (n = k) and then to all vertices belonging to the cluster with the closest centroid (n = V/k), resulting in a matrix with D_k_ elements:

$D_{k}=2\times S\times(k+\frac{V}{k})$	(C.2)

D_k_ is smaller than D if V is approximately smaller than k:

*D* < *D_k_*

$2\times S\times V$ < $2\times S\times(k+\frac{V}{k})$

$V<\frac{k^{2}}{k-1}\approx k$	(C.3)

By calculating the local minimum, it is easy to show that D_k_ assumes the smallest value at $k=\sqrt{V}$. Thus, using K-means with the number of clusters $k=\sqrt{V}$ reduces D by a factor of $\frac{D}{D_{k}}=\frac{V}{\sqrt{V}+\frac{V}{\sqrt{V}}}=\frac{V}{\frac{V+V}{\sqrt{V}}}=0.5\sqrt{V}=250$ at the typical value of V = 250000. Computational efficiency will be roughly proportional to D/D_k_ because, of the two operation types used in this filter (generation of the distance matrix and finding the smallest value within the matrix), the former is far more time-consuming.

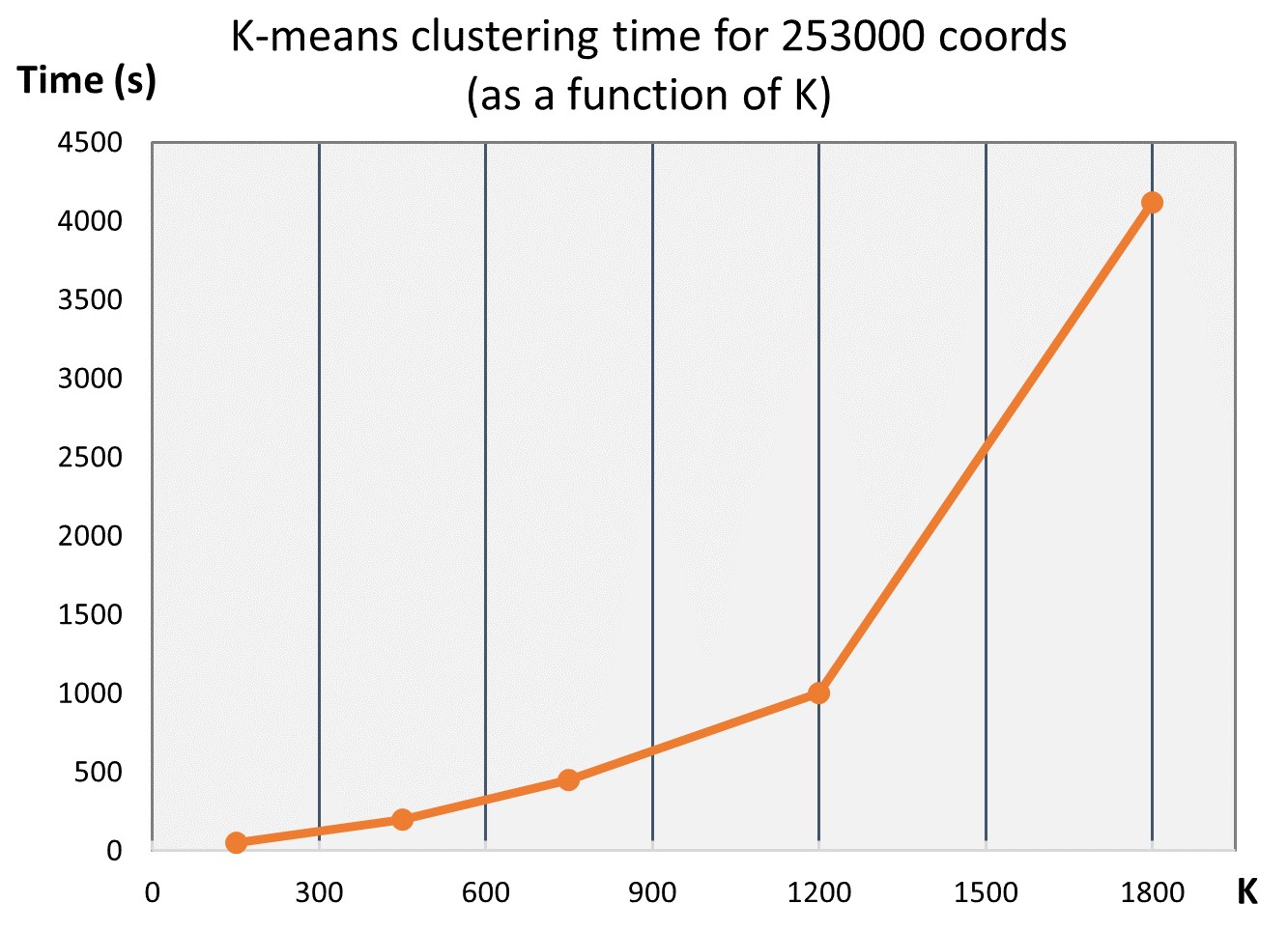

It is important to point out that K-means clustering will itself take an exponential amount of time depending on the number of clusters, with the typical V = 250000 and k = 500 resulting in about 4 mins during testing.

If a small number of streamlines is used (e.g., *S <* 10^5^), K-means clustering may actually prolong the overall GG filtering time. To avoid that, the framework will automatically set k to the arbitrary value of 60 when S is small, typically resulting in the whole process taking under 60 seconds during testing.
