## Appendix C for "Surface-based tracking for short association fibre tractography"

**Appendix C. The impact of the discrete voxel masks on filtering**

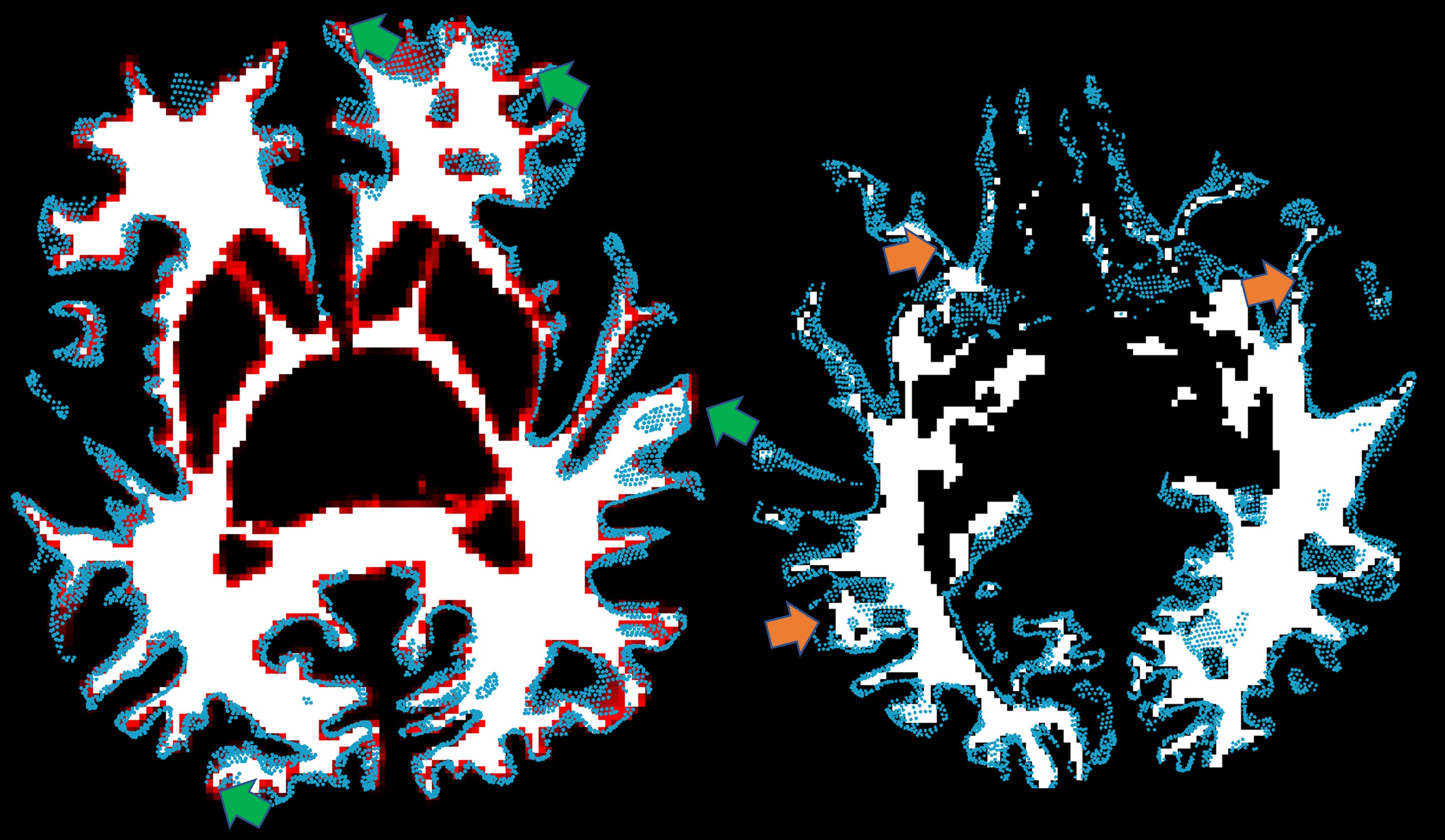

Figure C.1: Here the Freesurfer white matter mask was transformed from T1 to dMRI space using trilinear interpolation (producing antialiasing marked in red on the left-hand picture). WSM vertices are represented as blue dots (also in dMRI space). Some of the antialiased voxels reside outside the WSM (green arrows), which may cause problems with subsequent filtering. Removing the antialiaised voxels (right-hand picture, more inferior slice) will leave gaps (orange arrows) that will cause rejection of valid streamlines.
