## Appendix D for "Surface-based tracking for short association fibre tractography"

**Appendix D. Example of streamline data representation of the surface in a single subject**

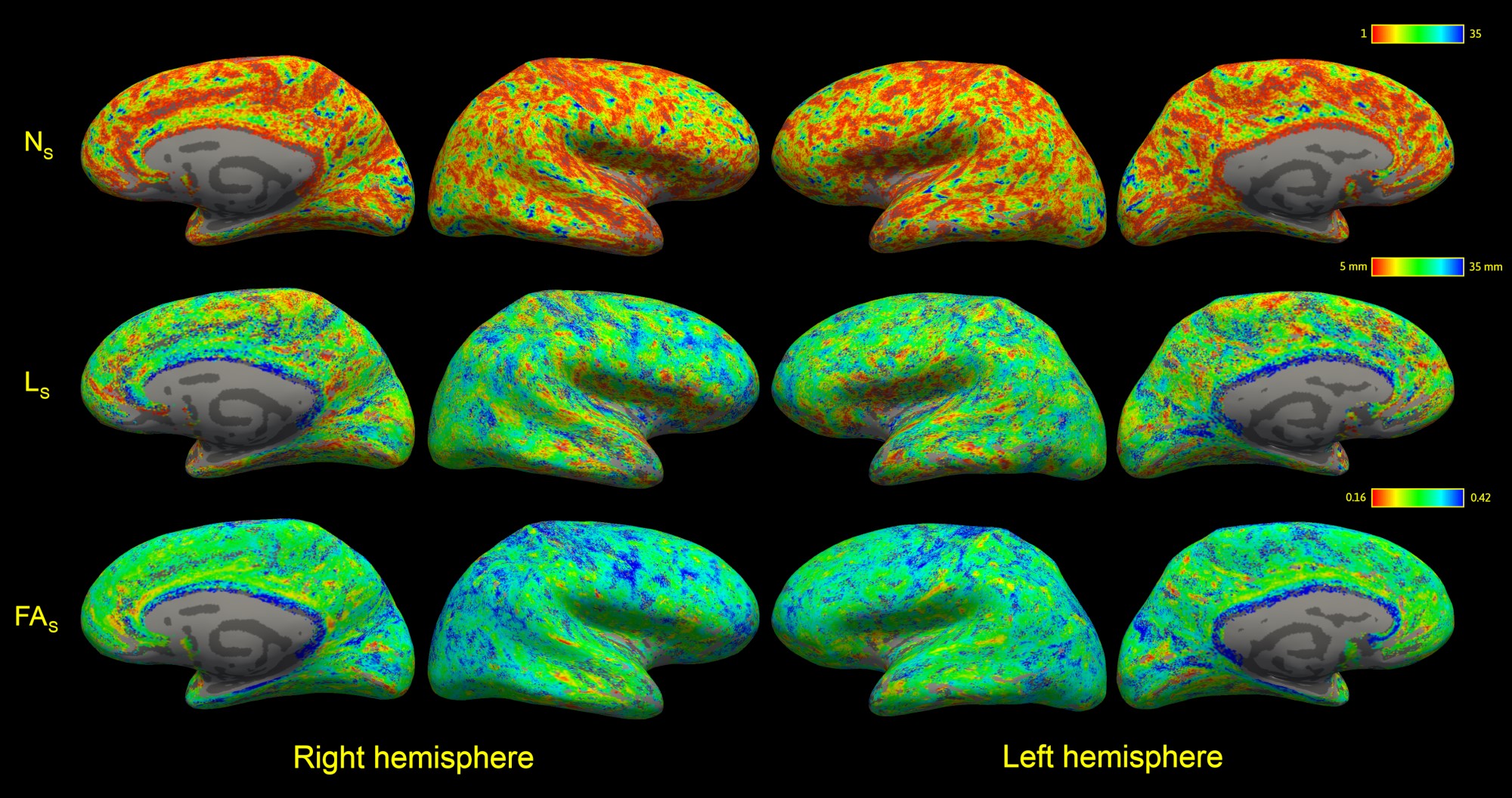

Figure D.1: Data shown are projected on the subject surface, with no smoothing applied. N_S_, termination density (number of streamlines/vertex). L_S_, mean streamline length/vertex. FA_S_, mean streamline fractional anisotropy/vertex.
