## Appendix E for "Surface-based tracking for short association fibre tractography"

### Appendix E. Misalignment between differently resampled data

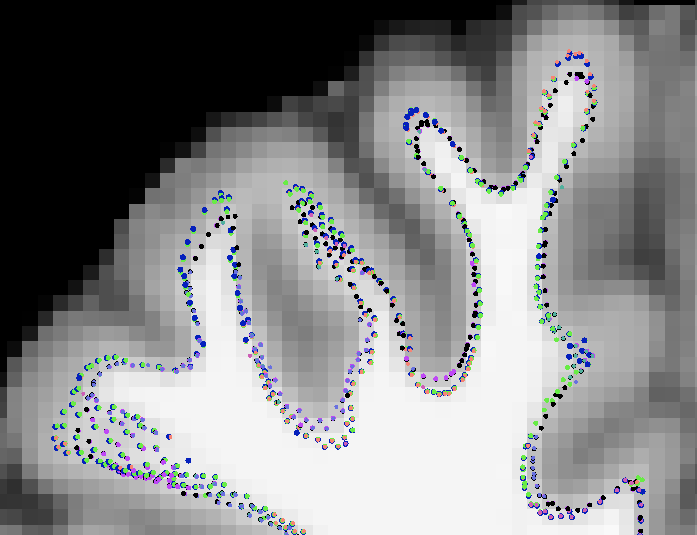
Figure E.1: Plot of white surface mesh vertices from all data sets together in the same DWI space after registration, overlaid on T1-weighted image. The same mesh coordinates were brought into the corresponding diffusion spaces (for surface-based tractography) which were then brought into the space of Prisma/SA/2mm isotropic data set (shown here) using FA volume co-registration. Most misalignment occurs in sulcal fundi and gyral crowns. These imperfections are exclusively between data sets with different voxel sizes: black, purple, lilac, cyan points show seeds from the 1×1×1 mm^3^ data while navy blue, light green, rose, pink (barely seen) points show seeds from the 2×2×2 mm^3^ data. The sulcal wall demonstrating the most difference in Figure 4 (yellow arrow) appears reasonably aligned.
