## Appendix F for "Surface-based tracking for short association fibre tractography"

**Appendix F. Consistency of SAF demonstrated using surface-based analysis**

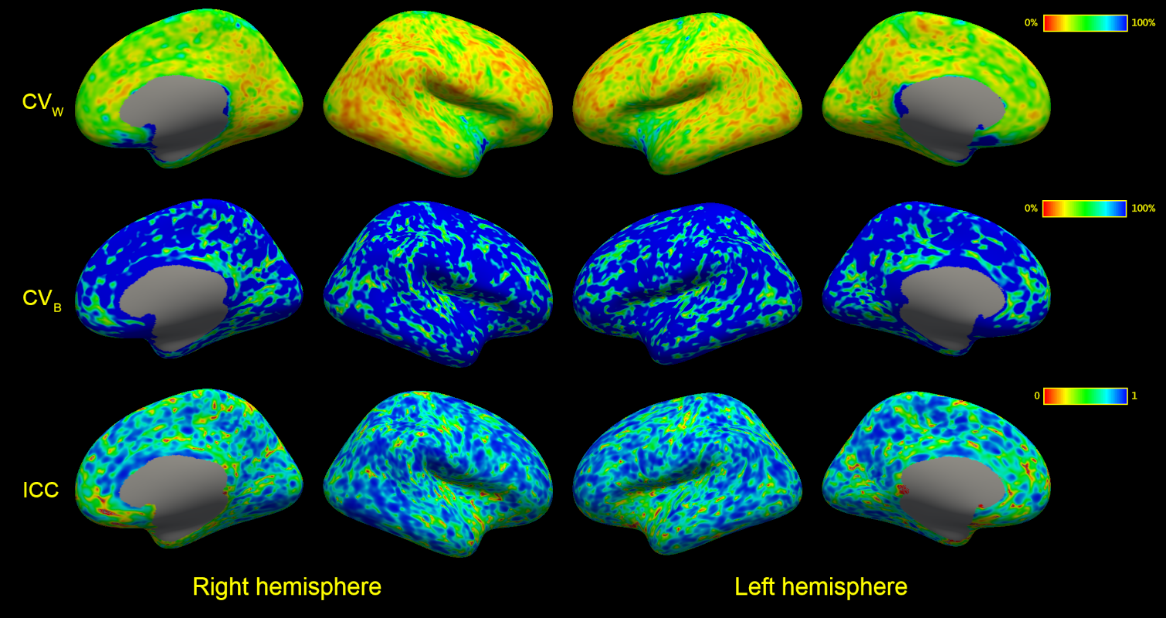

Figure F.1: Number of streamlines per vertex. CV_W_, coefficient of variation within subjects. CV_B_, coefficient of variation between subjects. ICC, intraclass correlation coefficient. Values at each vertex were recorded in subject space, then transformed into average subject space before running analyses. CV_W_ and CV_B_ were thresholded at 100%.

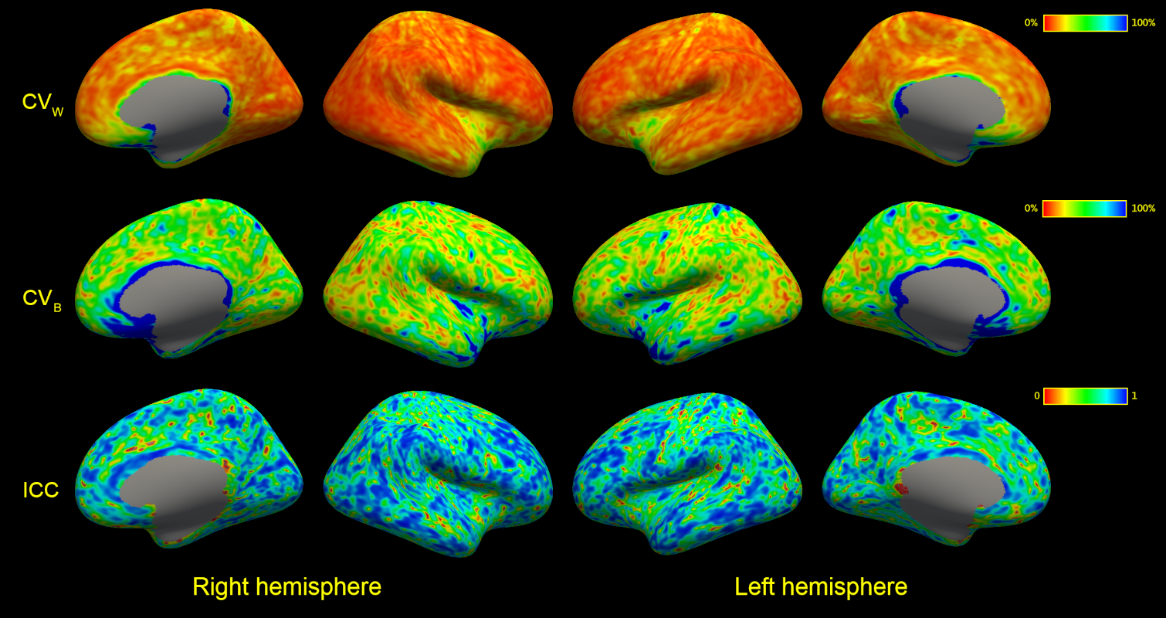

Figure F.2: Mean length of streamlines per vertex. CV_W_, coefficient of variation within subjects. CV_B_, coefficient of variation between subjects. ICC, intraclass correlation coefficient. Values at each vertex were recorded in subject space, then transformed into average subject space before running analyses. CV_W_ and CV_B_ were thresholded at 100%.

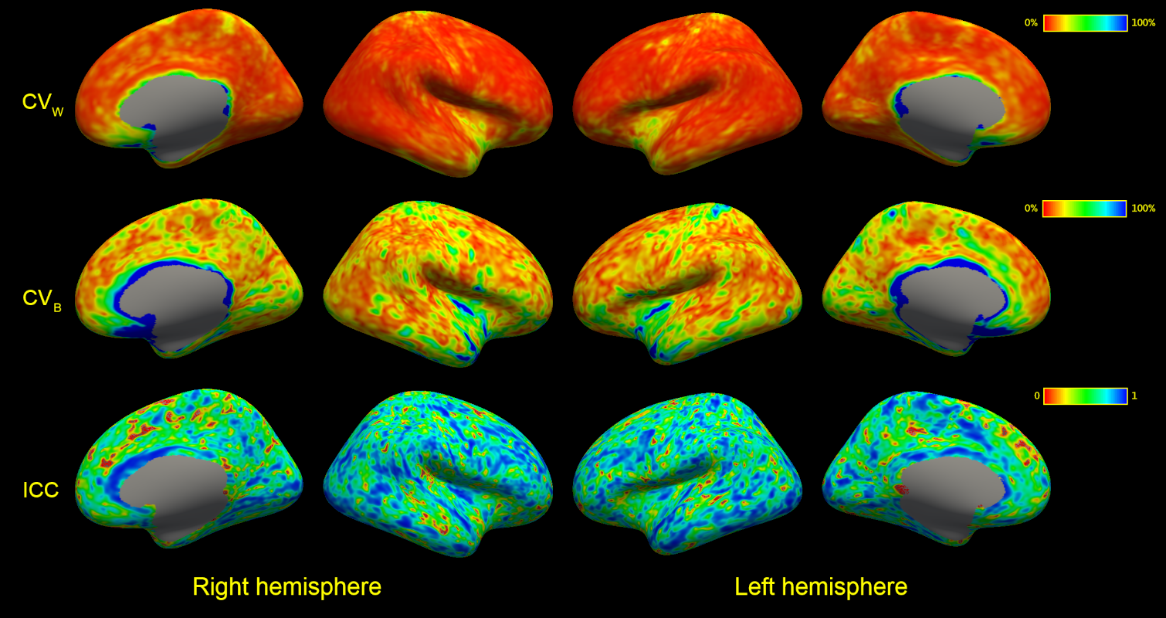

Figure F.3: Mean fractional anisotropy along streamlines per vertex. CV_W_, coefficient of variation within subjects. CV_B_, coefficient of variation between subjects. ICC, intraclass correlation coefficient. Values at each vertex were recorded in subject space, then transformed into average subject space before running analyses. CV_W_ and CV_B_ were thresholded at 100%.
